# An ancient ecospecies of *Helicobacter pylori* found in Indigenous populations and animal adapted lineages

**DOI:** 10.1101/2023.04.28.538659

**Authors:** Elise Tourrette, Roberto C. Torres, Sarah L. Svensson, Takashi Matsumoto, Muhammad Miftahussurur, Kartika Afrida Fauzia, Ricky Indra Alfaray, Ratha-Korn Vilaichone, Vo Phuoc Tuan, HelicobacterGenomicsConsortium, Difei Wang, Abbas Yadegar, Lisa M. Olsson, Zhemin Zhou, Yoshio Yamaoka, Kaisa Thorell, Daniel Falush

## Abstract

The colonization of our stomachs by *Helicobacter pylori* is believed to predate the oldest splits between extant human populations. We identify a “Hardy” ecospecies of *H. pylori* associated with indigenous groups, isolated from people in Siberia, Canada, USA and Chile. The ecospecies shares the ancestry of “Ubiquitous” *H. pylori* from the same geographical region in most of the genome but has nearly fixed SNP differences in 100 genes, many of which encode outer membrane proteins and host interaction factors. For these parts of the genome, the ecospecies has a separate, independently evolving gene pool with a distinct evolutionary history. *H. acinonychis*, found in big cats, and a newly identified primate-associated lineage both belong to the Hardy ecospecies and both represent human to animal host jumps. Most strains from the ecospecies encode an additional iron-dependent urease that is shared by *Helicobacter* from carnivorous hosts, as well as a tandem duplication of *vacA*, encoding the vacuolating toxin. We conclude that *H. pylori* split into two highly distinct ecospecies in Africa and that both dispersed around the globe with humans, but the Hardy ecospecies has gone extinct in most parts of the world. Our analysis also pushes back the likely length of the association between *H. pylori* and humans.

## Main text

*Helicobacter pylori* disturbs the stomach lining during long-term colonization of its host ^1,2^. Many strains harbour the *cag* pathogenicity island, which encodes a Type IV secretion system that injects a toxin, CagA, into host cells, interfering with cellular signalling and changing their morphology ^3^. Individuals infected by *cag^+^* strains have been found to have a higher disease burden of gastric cancer than individuals who are uninfected or infected by *cag^-^* strains ^4^. A second key host interaction factor, conserved in all *H. pylori*, is VacA, a vacuolating cytotoxin that shows substantial geographical variation ^1^. The factors maintaining this variation are unknown but could for example reflect dietary variation between human populations that necessitated different bacterial strategies for acquiring nutrients. If we could characterize these bacterial strategies, we would be better positioned to understand the nature of interactions between environment, host and bacteria that lead to the wide range of clinical outcomes in infected individuals ^2^.

We assembled a dataset of 6870 *H. pylori* and two *H. acinonychis* genome sequences from humans and other hosts around the world (Table S1, S2). *H. acinonychis*, has been isolated from big cats in zoos and represents a human-to-animal host jump ^5^. *H. pylori* have also been occasionally isolated from animals, including four from primates at the UC Davis Primate Research Center, which has housed rhesus macaques and Cynomolgus monkeys. In addition to published genomes ^6–32^, the dataset contains 2663 unpublished genomes from 54 countries, including a large new set of samples from Southeast Asia, and Iran, as well as strains previously only defined by Multilocus Sequence Typing, among them a high number of genomes from Siberia. Following previous practice (^25,26^), chromosome painting was used to assign the strains to 13 populations (designated hp) and less differentiated subpopulations (designated hsp), each of which have different geographic distributions (Table S2, Figure S1).

We noticed that alternative methods for clustering strains gave surprisingly different answers (Figure S2A, Figure S3, Figure S4). A subset of 48 “Hardy” strains from Chile, Siberia, Canada and the US as well as two strains from the Davis Primate Center formed a clade in phylogenetic trees (Figure S2A). The same strains were separated from the “Ubiquitous” others on a Principal Components Analysis (PCA) plot (Figure S3). Hardy strains were named as such because they were isolated from individuals living in locations that most of the world’s population would consider physically inhospitable, while “Ubiquitous” strains are found in humans everywhere. The human Hardy strains were assigned to three different populations, hspSiberia, hspIndigenousNAmerica and hspIndigenousSAmerica, and fineSTRUCTURE ^33^ grouped them with strains from their own geographic regions (Figure S4).

To understand why strains from different locations and ancestry profiles were grouped together by phylogenetic analysis and PCA, we performed a genome-wide analysis (GWAS, genome-wide association study) of differentiation between Hardy clade strains and other strains from the same fineSTRUCTURE populations. The differentiation was localized to specific genomic regions and was often confined by gene boundaries (Figure 1A, Figure S5). We therefore split the genome alignment into genes with nearly fixed differences between the Hardy clade and the Ubiquitous clade (100/1577), genes with no differentiated SNPs (1034) and intermediate genes that did not fall into the other two categories (443) to characterize patterns of genetic differentiation separately for these fractions of the genome (Table S3).

**Figure 1.**
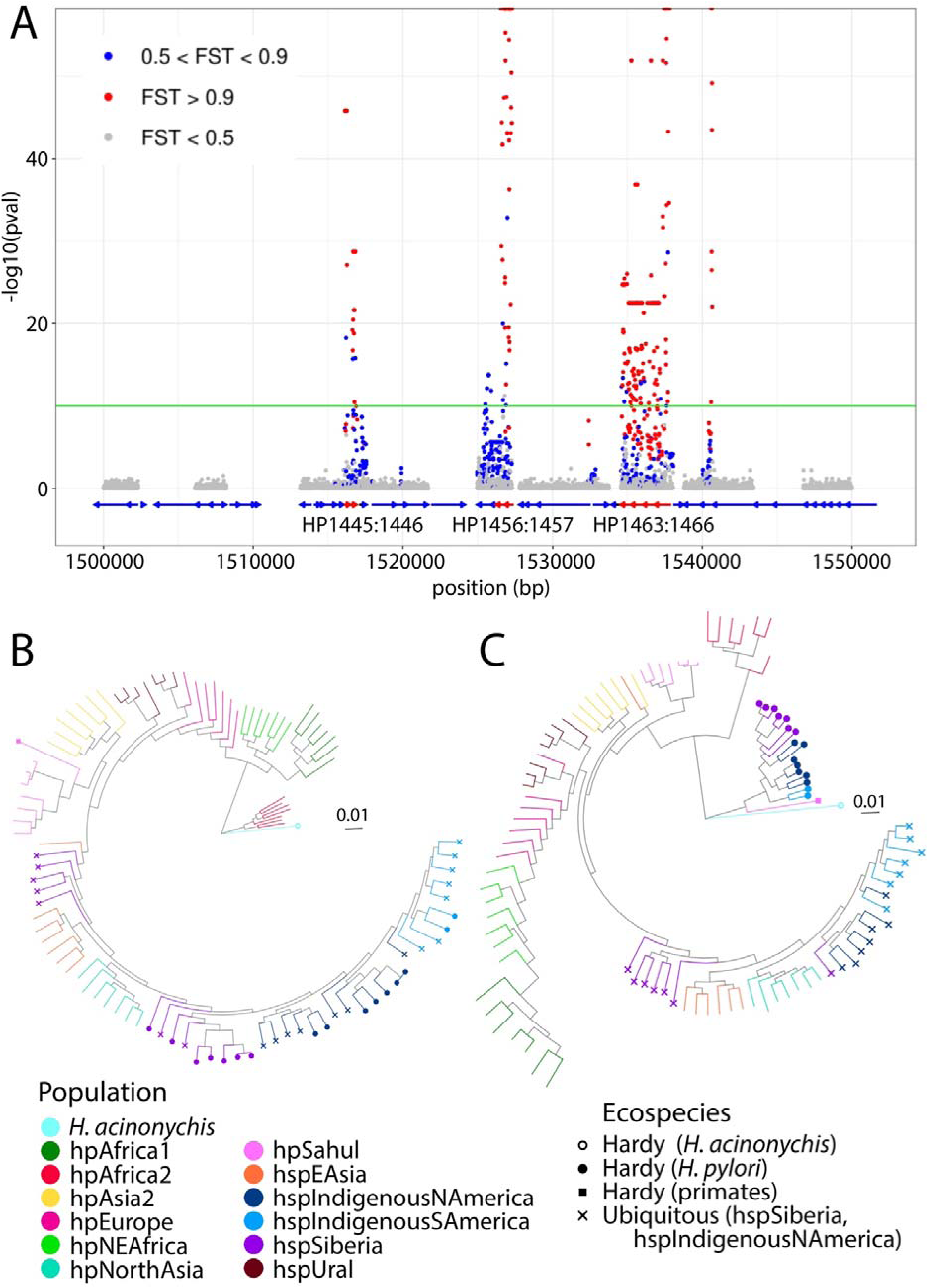
Differentiation between Hardy and Ubiquitous strains is localized in the genome. (A) Manhattan plot from GWAS analysis of the Hardy vs. Ubiquitous strains from hspSiberia and hspIndigenousNAmerica (zoom in between 1.5 and 1.55 Mbp, full plot shown in Supplementary Figure S5). Genes are indicated by blue and red (differentiated genes) arrows. Green line: significance threshold (-log10(p) = 10). Points are coloured based on FST (fixation index) values (red: FST > 0.9, blue: FST 0.5 - 0.9, grey: FST < 0.5). Half points at the top of the plot indicate an estimated p-value of zero and FST of one. Phylogenetic trees for undifferentiated genes (B) and differentiated genes (C) for a representative subset of strains (see Supplementary Figures S2B and C for trees of whole dataset). Branches are colored based on the population. Strains from the Hardy clade are indicated with a filled circle at the end of the branch.

Hardy strains have entirely different genetic relationships with other *H. pylori* at differentiated vs. undifferentiated genes, based on phylogenetic trees (Figure 1B, C, Figure S2B, C). The tree of undifferentiated genes (Figure 1B, Figure S2B) is consistent with the evolutionary relationships within *H. pylori* established in previous analyses ^16,34,35^, with Hardy strains from hspIndigenousSAmerica, hspSiberia and hspIndigenousNAmerica populations clustering with Ubiquitous strains isolated from the same locations. The longest branch in the tree separates strains from the hpAfrica2 population and *H. acinonychis* from other *H. pylori*. Concordant with previous inferences ^5,35^, the tree is rooted on this branch. hpAfrica2 strains originated in Khoisan populations in Southern Africa ^35^, who are the oldest-branching group in the human population tree ^36^. At differentiated genes (Figure 1C, Figure S2C), the Hardy strains isolated from humans are the most genetically distinct *H. pylori*, branching more deeply than hpAfrica2 and are more closely related to *H. acinonychis*. This divergence causes the Hardy strains to branch separately from Ubiquitous strains from the same population in phylogenetic analyses based on the whole genome (Figure S2A).

The two primate Hardy isolates are assigned to the hpSahul population, found in humans in Indonesia, Papua New Guinea, and Australia and cluster with other isolates from this population in the tree of undifferentiated genes (Figure 1B, Figure S2B). However, the ancestry profile of these isolates is clearly distinct from those of any other hpSahul isolate in the database (Figure S6), suggesting the host jump may have been ancient. Two further strains from the UC Davis Primate Research Center belong to the hpAsia2 population (Table S2) and appear to be typical Ubiquitous *H. pylori* (data not shown).

We next investigated the clonal relationships of Hardy and Ubiquitous. Bacteria reproduce by binary fission, meaning that there is a single genealogical tree representing the clonal (cellular) relationships of any sample, but recombination can also transfer DNA between strains. Recombination of short tracts is unlikely likely to affect genome order, especially for large rearrangements. Therefore, rearrangements can potentially provide a marker of clonal descent, even in the presence of high recombination rates. Dot plots revealed many more rearrangements between Hardy and Ubiquitous strains than between strains of the same type, even when the Hardy and Ubiquitous strains came from the same population (Figure S7). These results show that at the cellular level, the split between Hardy and Ubiquitous strains was ancient. The presence of around 140 rearrangements between strains from the same location ^37^ has done little to suppress gene flow in most of the genome due to the short length of many fragments incorporated by homologous recombination in *H. pylori* ^38^. Nevertheless, at the differentiated genes, Hardy and Ubiquitous strains have kept their distinct identities.

To reconcile the discordant evolutionary history of differentiated and undifferentiated genes and the ancient split between the Hardy and Ubiquitous clonal backgrounds, we hypothesize that *H. pylori* split into two ecospecies prior to the split of hpAfrica2 from other populations (Figure 2A). We define ecospecies as bacteria within the same species that have undergone species-level differentiation within a specific fraction of the genome, while otherwise having a single recombining gene pool. It is likely that differentiation in that fraction is maintained by strong ecologically mediated selection against intermediate genotypes, but other forces, such as genetic mechanisms greatly restricting gene-flow in parts of the genome might also play a role.

**Figure 2.**
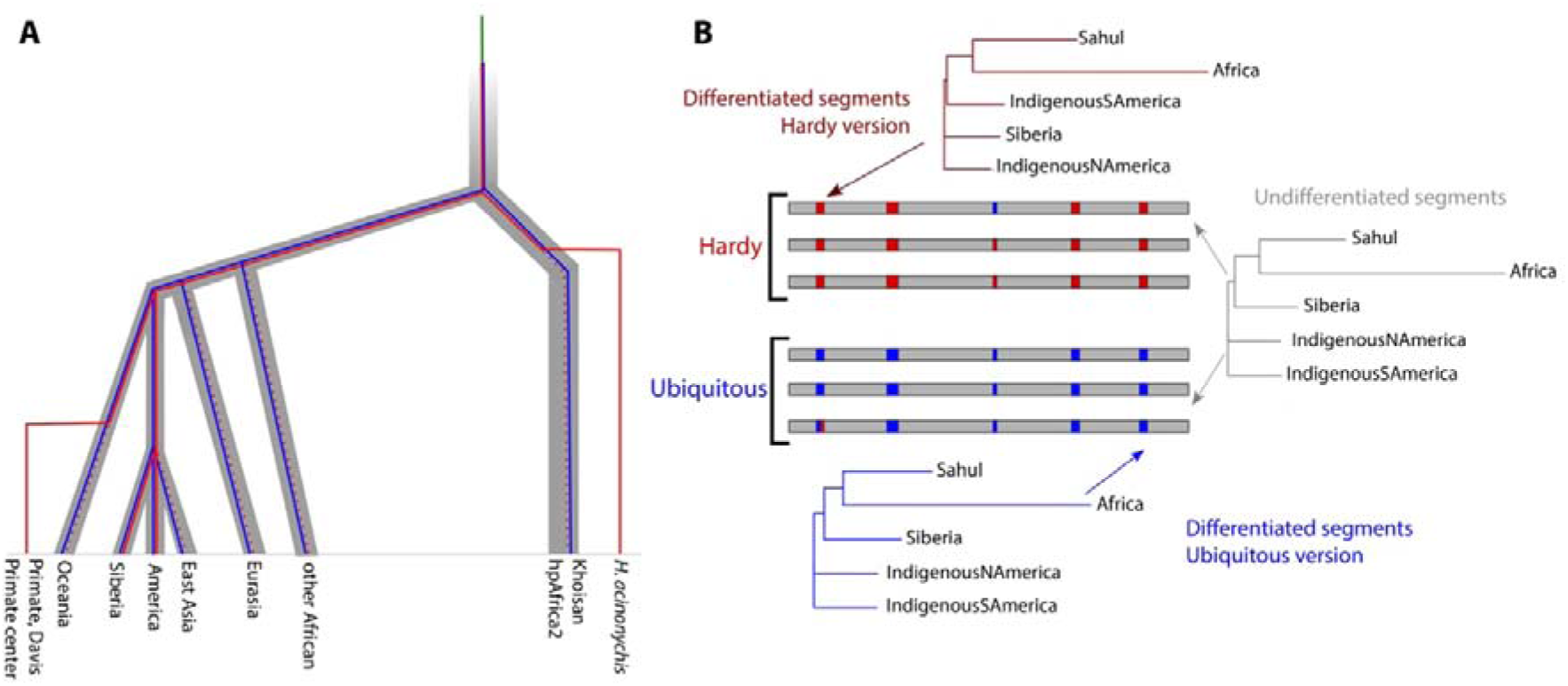
Divergence and spread of Hardy and Ubiquitous gene pools. (A) Hypothesized scenario for the differentiation of *H. pylori* into two ecospecies and subsequent global spread. Thick grey tree: simplified history of human population differentiation (based on ^40^). *Helicobacter* evolution is represented by thinner lines, which are within the grey tree during periods of evolution in humans. Green line: *H. pylori* prior to the evolution of the two ecospecies. Blue line: Ubiquitous ecospecies. Red line: Hardy ecospecies. Dotted red lines indicate that Hardy strains have not yet been detected on the branch and therefore may have gone extinct. Not to scale. (B) Phylogenetic population trees for the differentiated (Hardy in red and Ubiquitous in blue) and undifferentiated (in grey) regions of the genomes. Trees were constructed considering only populations with both Hardy and Ubiquitous representatives. Pairwise distances between strains were calculated for the relevant genome regions, and population distances were calculated by averaging over pairwise strain distances. For Africa, we used *H. acinonychis* strains for the Hardy differentiated gene trees and hpAfrica2 strains for the Ubiquitous differentiated genes tree. For the undifferentiated gene tree, we averaged over the relevant Hardy and Ubiquitous strains.

We propose that both ecospecies were spread around the world by human migrations. Since the gene pools at differentiated loci are distinct, with their own segregating SNPs, genetic relationships at these genes represent the separate histories of spread of each ecospecies. The DNA may have been carried to their present locations by different human migrations, while undifferentiated genes reflect an amalgam of both histories. Phylogenetic trees constructed for Hardy and Ubiquitous strains from the same populations at differentiated genes show similar, parallel, histories (Figure 2B), albeit with minor differences. Specifically, for Ubiquitous and undifferentiated segments, hspIndigenousSAmerica and hspIndigenousNAmerica strains cluster together. In Hardy strains, hspIndigenousSAmerica strains cluster in a deeper position in the tree, hinting that they may have been brought to the Americas by a distinct, and probably earlier, human migration.

To estimate the age of the ecospecies, we need a consistent molecular clock. Synonymous divergence levels, dS, are similar for Ubiquitous strains in differentiated and undifferentiated genes (Figure S8A and B), implying no large difference in mutation rates. In differentiated genes, comparisons between ecospecies gave divergence levels greater than 0.3 for all comparisons involving *H. acinonychis* or hpAfrica2 as outgroups (Figure S8B and D – *H. acinonychis* subplot). Comparisons between hpAfrica2 and other Ubiquitous strains gave consistently lower dS values (average dS = 0.23, Figure S8B), implying that the ecospecies arose substantially before the divergence of hpAfrica2 from other *H. pylori.* Comparisons involving human Hardy strains showed intermediate dS values (Fig S8D, Hardy *H. pylori* subplot), with the lowest values for strains where Hardy and Ubiquitous coexist. We attribute this to residual gene flow even within the differentiated regions, which, as expected, is weaker at non-synonymous sites. This gene flow will cause us to systematically under-estimate divergence times.

Previous analysis ^35^ suggested that hpAfrica2 split from other *H. pylori* around 100,000 years ago and concluded that there is no direct evidence for *H. pylori* in humans before that. Based on dS values between hpAfrica2 and *H. pylori* Hardy strains (Fig S8B and D – Hardy *H. pylori* subplot) our analysis implies that Hardy and Ubiquitous evolved at least 0.34/0.23 before this split, or at least 150,000 years ago. However, an alternative hypothesis is that hpAfrica2 strains codiverged with Khoisans, which according to current human genetic models are the oldest branching extant human group, with a split time from other human populations of around 200,000 years, albeit with considerable uncertainty ^36,39^. If this is instead used as a calibration point for Hardy evolution, then this minimal date estimate is pushed back by a factor of 2 to 300,000 years.

The ratio of non-synonomous to synonymous diversity, dN/dS, indicates the degree of functional constraint on genes and are also impacted by positive selection (Figure S8A and B). Within Ubiquitous *H. pylori*, dN/dS levels are lower in differentiated genes than in undifferentiated ones (0.125 vs 0.171), implying than differentiated genes are on average more functionally constrained than the rest of the genome. However, at differentiated genes, dN/dS levels are elevated in the pairwise comparisons involving Hardy and hpAfrica2 strains (0.178), which implies that after the differentiation between the two ecospecies there was a burst of evolution at functional sites at these genes, which is likely associated with their specialization to distinct niches.

The fineSTRUCTURE results, with similar levels of coancestry within and between ecospecies in particular geographical locations, demonstrate that there is frequent recombination between ecospecies in most of the genome (Figure S4). The question is then what force has maintained the differentiated regions? If this force is natural selection, then we should see genetic exchange between ecospecies that is too recent to have been purged. The distribution of haplotypes in the differentiated regions fits with the action of natural selection (Figure 3A), with sharing of segments between the Hardy ecospecies and other strains in many genes but with low frequencies for each imported fragmen.

**Figure 3.**
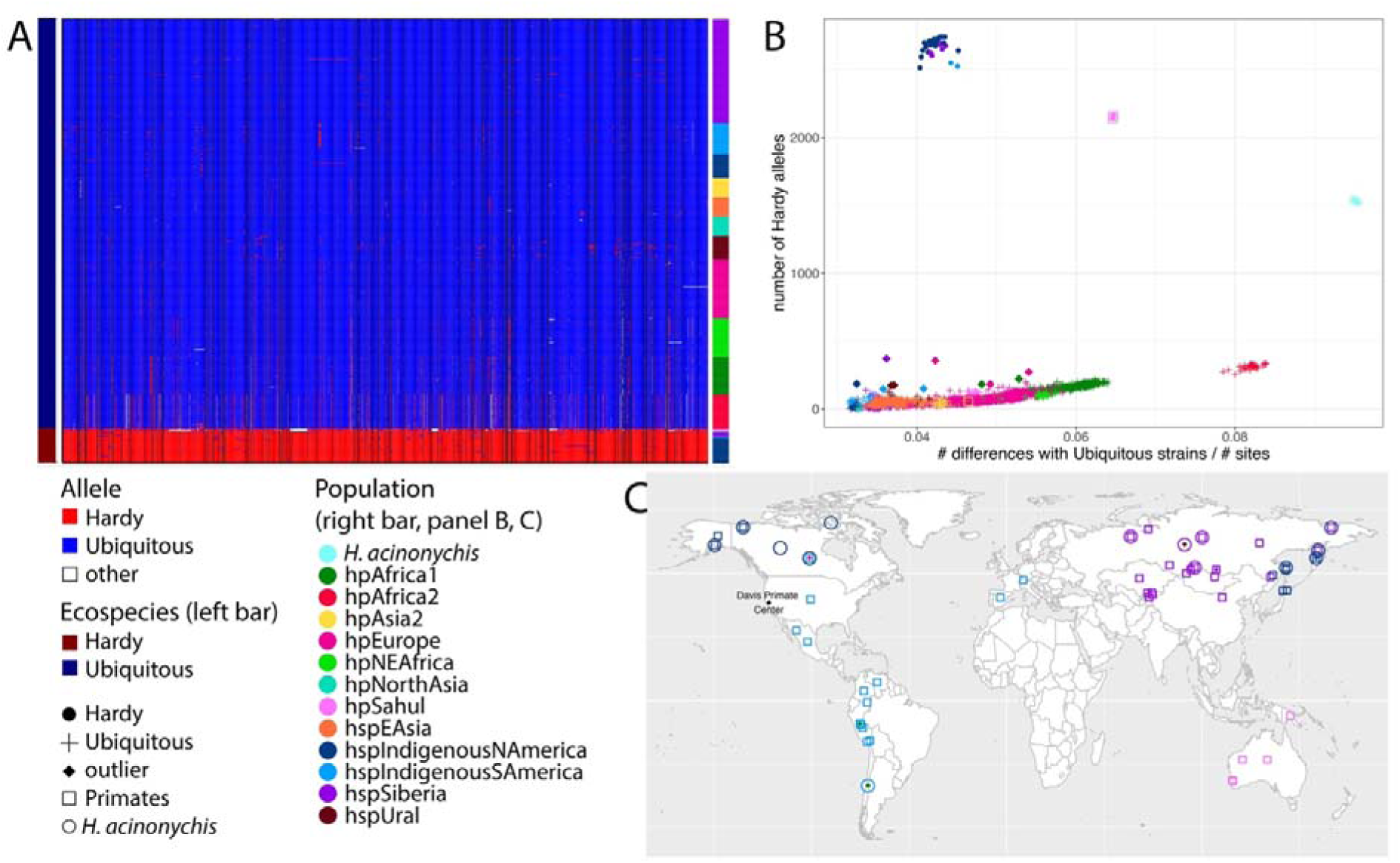
Geographic distribution of Hardy and Ubiquitous haplotypes. (A) Haplotypes at the SNPs that are differentiated between the two ecospecies for hspSiberia and hspIndigenousNAmerica strains and randomly selected representatives from other populations. The major Hardy allele is represented in red while the major Ubiquitous allele is in blue. Other alleles are shown in white. The black vertical lines separate the different genes. (B) Number of Hardy alleles as a function of average genetic distance with the Ubiquitous ecospecies strains from hspIndigenousSAmerica, hspSiberia and hspIndigenousNAmerica. The dots are coloured based on their populations and the Hardy strains are represented with a circle. Crosses indicate outlier strains with a higher number of Hardy alleles than expected for a strain at that genetic distance. The inset is a zoom around these outliers. (C) Map with the location of the Hardy (circle) and Ubiquitous (square) strains from hspIndigenousSAmerica, hspSiberia, hspIndigenousNAmerica and hpSahul populations as well as the strains that were outliers (full diamond) compared to their population in the number of Hardy alleles they have. For some of the strains, the information on where they were isolated (latitude and longitude) was missing. In this case, we put the coordinates of their country of isolation instead.

Further evidence for selection maintaining the distinction between ecospecies comes from the limited geographic range of haplotypes originating in the Hardy clade. To identify strains that have received such DNA, we plotted the number of Hardy alleles, i.e. the predominant allele in the Hardy clade, as a function of genetic distance to representatives of the Ubiquitous ecospecies from populations where the Hardy ecospecies strains were isolated. Distantly related strains (e.g. hpAfrica2) have a higher number of such alleles, but this can be attributed to differentiation of the representative Ubiquitous strains rather than to gene flow between ecospecies, as can be seen by the consistent upward trend in Figure 3B. There are 11 strains (Table S2) that are clear outliers from the trend line in Figure 3B, and all of these come from geographic regions where the Hardy ecospecies is found (Figure 3C). The same outliers appear in a plot of the number of differentiated blocks against the number of differentiated SNPs (Figure S9). Since genetic exchange tends to progressively homogenize ancestry in the absence of natural selection, the absence of outlier strains elsewhere implies that Hardy alleles are deleterious on a Ubiquitous genetic background and the outliers themselves can be explained by recent, local genetic exchange. Furthermore, the absence of outliers elsewhere implies that Hardy has itself has had a limited geographical range in recent times.

Our hypothesis presented in Figure 2A implies that Hardy progressively diverged from Ubiquitous via new mutations. To investigate evidence for alternative scenarios in which acquisition of DNA from other *Helicobacter* played a role, we assembled a panel of *Helicobacter* species from other mammals, excluding *H. acinonychis* (Table S4). We performed BLAST on each of the 100 differentiated genes to identify close matches. In total, 49 genes had significant BLAST matches (Table S3), the closest of which was always in *H. cetorum*, whose hosts are dolphins and whales and is the closest known relative of *H. pylori* other than *H. acinonychis*.

The results of the BLAST analysis show no evidence for DNA being imported to *H. pylori* from other *Helicobacter*. For 29 of the genes, the evolutionary distances, visualized using a phylogenetic tree (Figure S10A), were consistent with the scenario shown in Figure 2A, with Hardy and Ubiquitous versions diverging from each other before the split of hpAfrica2 / *H. acinonychis* from other populations. For 13 of the genes, the tree instead implied that the Hardy and Ubiquitous versions of the gene diverged from each other after the split between hpAfrica2 and other human *H. pylori*, with hpAfrica2 and *H. acinonychis* versions of the gene clustering together (Figure S10B). Three genes (*horL*, methyl-accepting chemotaxis protein HP0599, and *lp220*) showed signs of gene flow between Hardy and Ubiquitous versions (Figure S10C). Three genes, including *vacA* (discussed below) had a history involving gene duplication (Figure S10D), while for the gene encoding the outer membrane protein HopF, the Hardy version diverged from the Ubiquitous version before the split with *H. cetorum* (Figure S10E). This pattern could be explained by cross species genetic exchange, but an alternative is that the common ancestor of *H. pylori* and *H. cetorum* had two different versions of the gene, with both copies being maintained in *H. pylori* until it split into ecospecies. There was no other gene where the pattern of polymorphism was suggestive of import of genetic material from other *Helicobacter* species.

Clues to the ecological basis of ecospecies divergence are provided by the functions of differentiated and accessory genes from the Hardy ecospecies (Table S3) Hardy strains, including *H. acinonychis* form their own clade in a tree constructed based on accessory genome analysis (Figure 4A). Only five (the two primate strains included) out of 48 Hardy strains have an intact *cag* pathogenicity island, which is a much lower rate than for the Ubiquitous ecospecies (472/673 strains), but enough to imply that the island can function and be maintained by natural selection on both ecospecies backgrounds (Figure 4A and B). The *vacA* gene, coding for vacuolating cytotoxin, is one of the 100 genes differentiated between ecospecies and in addition, the completely sequenced Hardy genomes reveal an additional tandem pair of duplicated *vacA* genes (Figure 4B and C, Figures S10, S11).

**Figure 4.**
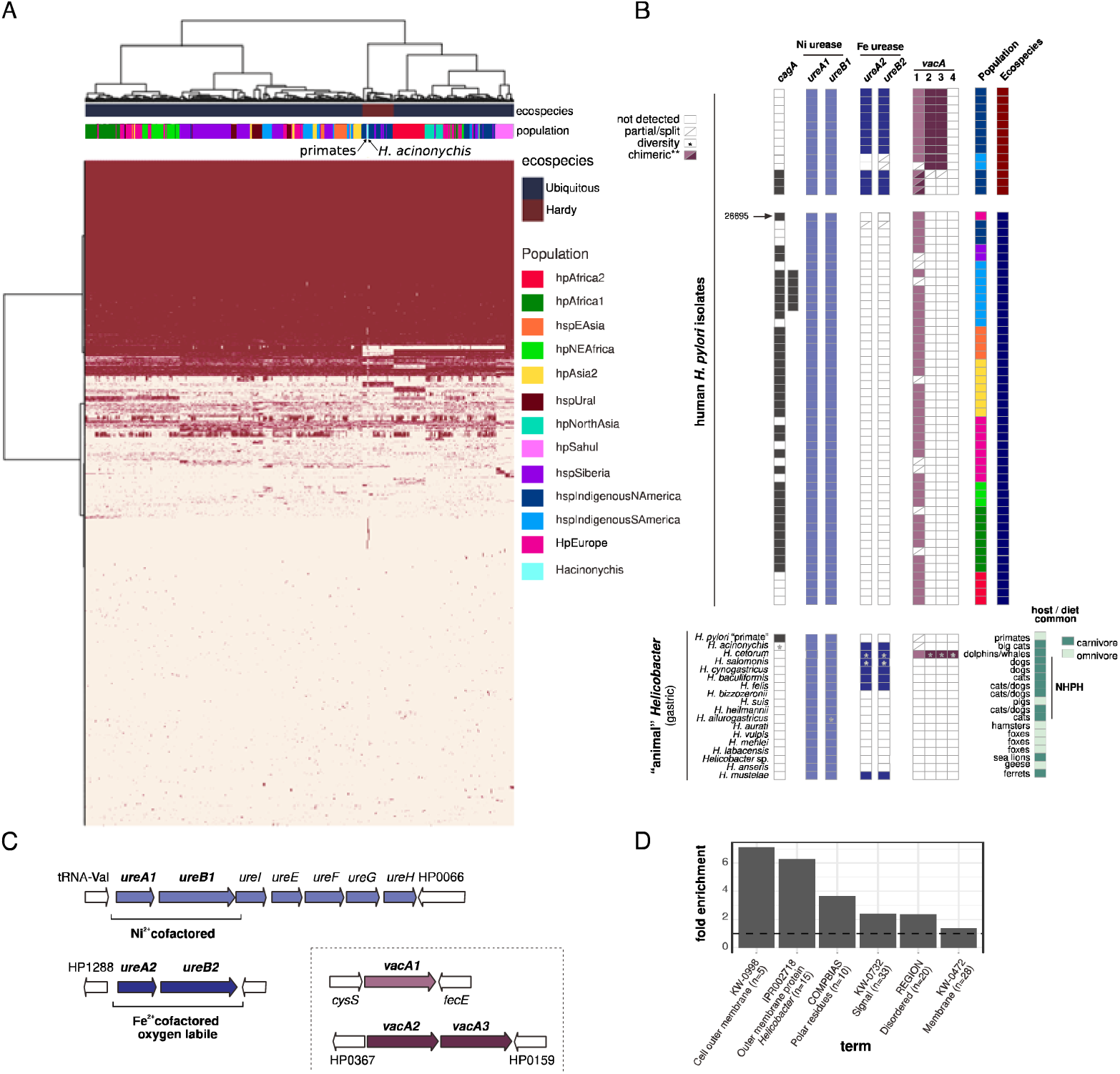
Genome composition in human and animal *Helicobacter*. (A) Pangenome analysis on a sample of the global dataset. Red: presence, White: absence. Strains are colored based on their population and their ecospecies. (B) (Top) Presence/absence of *cagA* and different *vacA* and *ureA/B* types in Hardy and Ubiquitous *H. pylori* from humans. Included is a subset of complete genomes, two *cagA*+ Hardy strains, and the reference strain 26695. (Bottom) Presence/absence in gastric *Helicobacter* isolated from animals. Common host(s) and their diets are indicated. Diversity: differential presence/absence in analyzed genomes within the species. Chimeric: potential chimeric version (Hardy + Ubiquitous). NHPH: non-*H. pylori Helicobacter* species. See Table S4 for details. (C) Representative configuration and genomic context of *ureA/B* and *vacA* in Hardy *H. pylori* genomes. Based on strain HpGP-CAN-006. Lighter coloured arrows: Ubiquitous versions, based on sequence and genomic context/synteny. Darker colours: Hardy versions. (D) Fold enrichment for significant (p < 0.05, Fisher’s test, Benjamini correction) functional terms.

Hardy strains carry an iron-cofactored urease (UreA2B2) in addition to the canonical nickel-dependent enzyme of *H. pylori* (Figure 4B and C). Iron-cofactored urease is thought to allow survival in the low nickel gastric environment of carnivores ^41^. Although previously noted in a single Hardy *H. pylori* strain from Northern Canada ^37^ this feature has otherwise been entirely associated with *Helicobacter* residing in obligate carnivore animal species: *H. mustelae* (ferrets), *H. felis*/*H. acinonychis* (cats, big cats), and *H. cetorum* (dolphins, whales) ^42^. Pangenome analysis of non-*pylori Helicobacter* species isolated from animals (Figure 4B, Table S4) confirmed these associations, and also identified *ureA2B2* homologs in additional species associated with carnivores (cats, dogs) such as *H. cynogastricus*, *H. salomonis*, and *H. baculoformis*. We did not detect *ureA2B2* in some gastric *Helicobacter* associated with cats and dogs (e.g., *H. heilmannii*, *H. bizzozeronii*); however, these species are known to co-colonize the stomach with other species that carry the iron-urease such as *H. felis* ^43^. We did not detect *ureA2B2* in any species that colonize omnivores, in the Hardy primate strains that also likely encounter a varied diet (Figure 4C), or any enterohepatic species (data not shown). The *ureA2B2* locus reflects a very old duplication event, with versions of both genes also found in *H. cetorum* (Figures S12, S13). Therefore, as for *vacA*, it is likely that the ancestral population had two versions of the gene, but that one has been lost by Ubiquitous strains after the evolution of the ecospecies.

Functional enrichment analysis of the 100 differentiated genes in the Hardy ecospecies confirmed a striking enrichment of *Helicobacter* outer membrane proteins (21/100 genes, p<0.05, Figure 4D, Table S3, S5) and the list also included many genes involved in cell envelope biogenesis. Four Outer Membrane Protein coding genes are unique to the Hardy genomes. One gene resembles the carbohydrate-binding adhesin BabA, which is missing in the Hardy strains, but major alterations in the cysteine-loop forming motifs involved in glycan binding suggest differences in binding specificity.

## Discussion

Our analysis has found that Indigenous people in Siberia and North and South America are infected by two distinct types of *H. pylori* that we have named Hardy and Ubiquitous. These types are distinguished by nearly fixed differences in 100 out of 1577 genes (based on an alignment with the 26695 reference genome) and important differences in gene content, despite evidence of extensive genetic exchange throughout the genome. We designate these types as ecospecies, since within the differentiated fraction of the genome they have their own gene pools which are sufficiently different (dS = 0.34), that they would be considered a different species based on usual criteria applied to bacteria ^44^, if this divergence was found across the genome. However, they are not species, since most of the genome is undifferentiated and there is no evidence that distinct types ever existed in the undifferentiated fraction of the genome.

The existence of these highly distinct genotypes within multiple geographical regions poses the question of how they have arisen and spread. Our explanation emphasizes continuity within their established primary host species, namely humans. The two ecospecies currently coexist stably within several human populations and we propose that this has been continuous since the distinction between the types first arose in the ancestor of modern humans. This hypothesis can explain the current pattern of diversity parsimoniously, without requiring either host jumps ^37^ or convergent evolution ^45^.

Our genomic data provides evidence against plausible introgression hypotheses for the two ecospecies. For example, if humans came out of Africa carrying Ubiquitous *H. pylori* and these hybridized with bacteria with Hardy alleles that had evolved in Neanderthals, Denisovans, big cats or aquatic mammals, then this hybridization should have generated genetic exchange throughout the genome, consistent with high inter-population recombination rates observed wherever distinct *H. pylori* populations coexist. However, in the non-differentiated fraction of the genome, we see no evidence for elongated branch lengths in those populations where the Hardy ecospecies is found (Figure 1B, Figure S2B), that would be generated by this gene flow, nor evidence of an elevated genetic distance to hpAfrica2 (Figure S8A).

A second argument against a hybrid origins hypothesis is that the 100 genes that define Hardy and Ubiquitous ecospecies have an approximately parallel history of spread, which takes in Africa, Sahul, Siberia, North and South America, as can be seen in the similar topology and branch lengths of phylogenetic trees for these loci (Figure 2B). A simple explanation for this approximately parallel history is that both types have been spread to these continents by the same or similar migrations of modern humans, before subsequent host jumps to big cats (*H. acinonychis*) and primates (Davis Primate Center isolates) and transportation of animals by humans within historical times. If the historical reservoir for Hardy was an archaic human or an animal, it is hard to explain the parallel history of spread or indeed how these disparate locations were reached. Few species have been as successful in colonizing the world in the last 100,000 years as modern humans.

While there is no evidence of import of diverged DNA into human *H. pylori* populations, as required by an introgression hypothesis, there is good evidence of progressive divergence within humans at the loci that define the ecospecies. According to the gene-by-gene topologies presented in Figure S10, for around 1/3 of the ecospecies defining genes, the divergence between Hardy and Ubiquitous versions commenced after the split from hpAfrica2 population, in other words within the history of extant human populations. Furthermore, there is evidence that positive selection has precipitated divergence, in the form of elevated dN/dS levels in comparison between ecospecies, at ecospecies defining genes (Figure S8D).

Evidence that distinct ecospecies can arise and be maintained within a single bacterial population is provided by the free living bacteria *Vibrio parahaemolyticus*, which alongside *H. pylori* is the most recombinogenic bacteria found to date ^46^. *V. parahaemolyticus* has a deeply diverged ecotype (EG1a), involving extensive coadaptation between loci encoding lateral flagella and genes involved in cell wall biosynthesis. This ecotype is also shared between geographical gene pools and has 18 diverged genome regions ^47^. Therefore, it also meets the ecospecies criteria and is also characterized by elevated rates of non-synonymous mutation at ecospecies defining genes. In *V. parahaemolyticus* there are also less differentiated types EG1b, EG1c and EG1d, which may represent precursors to ecospecies differentiation. Since both highly recombined species identified to date have clearly defined ecospecies, it is an open question how frequent they are within bacteria.

Notwithstanding the evidence for progressive evolution of Hardy and Ubiquitous ecospecies within humans, several important uncertainties remain. Our estimate is that the ecospecies are at least 1.5 times older than the split of hpAfrica2 from other *H. pylori* groups, but we have little evidence on the evolution that took place before that time. It is possible that initial divergence took place in distinct host populations and this initial divergence may have been non-adaptive. Furthermore, in the absence of functional assays, we cannot be sure what the selective forces are that have maintained the distinction between ecospecies and their long-term coexistence. To gain further insight into host-pathogen coevolution, it will be important to understand why the Hardy ecospecies has been able to reach Sahul, North and South America and Siberia from Africa but has now disappeared from most of the world.

One speculative factor that could explain ecospecies distribution is variation in human diets during our colonization history. Hardy strains share an iron-cofactored urease with *Helicobacter* from carnivores. In mammals, the main source of nickel, the cofactor of urease in Ubiquitous strains, is thought to be plant-based ^41^. This poses a challenge to *Helicobacter* colonizing the stomachs of carnivores. The iron urease is thought to allow persistent colonization by species such as *H. felis* and *H. mustelae* even when nickel is limiting. Further evidence for the potential role of diet is that several other genes that are differentiated between Hardy and Ubiquitous strains encode iron and nickel acquisition factors (*frpB4*, *tonB1*, *exbB*, *exbD*) or iron/acid adaptation regulators (*fur*, *arsS*).

Humans themselves have come under selection pressure due to changing diet. Hardy strains are currently found in human populations where few crops grow and that have ancestral alleles at the *FADS* (fatty acid desaturase) locus ^48^, responsible for fatty acid metabolism, that have been strongly selected against within European populations over the last 10,000 years ^49^. The modern distribution of *H. pylori* ecospecies could be explained if humans relied principally on hunting when colonizing new locations but this depleted large prey ^50^ and therefore that the first human migrations transmitted both ecospecies, but that Hardy died out in most locations after the initial colonization wave. If so, this analysis also implies that the ancestral population in which Hardy and Ubiquitous first differentiated was also heavily dependent on hunting. Functional work will be necessary to establish the conditions in which Hardy and Ubiquitous strains are able to coexist and to further flesh out these hypotheses.

There is an intimate relationship between *H. pylori* pathogenesis and host iron status, and nutritional adaptation has also been proposed as one explanation for the benefit of virulence factors such as VacA and CagA to the bacterium ^2^. The Hardy ecospecies has been isolated from human populations with significant gastric disease risk ^37^ and *H. acinonychis* causes severe gastritis and is a frequent cause of cheetah death in captivity ^51^. However, it remains to be seen if the Hardy ecospecies can also be defined as a "pathotype". In addition to genes associated with metal uptake, a large fraction of the genes that are differentiated between ecospecies are in outer membrane proteins, including adhesins, and other genes associated with virulence. This association is fascinating as it could shed light on the interaction between human diet, bacterial colonization strategies and virulence.

## Methods

### Genome collection

We collected a total of 9188 *Helicobacter* whole genome sequences from public and private sources, including 5493 *H. pylori* and 2 *H. acinonychis* genomes publicly available in Enterobase ^52^ (as of Jun 6th 2022), 1011 samples from the “*Helicobacter pylori* genome project” (HpGP) ^26^ (https://doi.org/10.5281/zenodo.10048320) and 286 samples available in NCBI, BIGs and FigShare; and 3635 *H. pylori* novel genomes from different geographic regions around the world. The novel sequences included 2126 isolates collected by Professor Y. Yamaoka at the Department of Environmental and Preventive Medicine, Faculty of Medicine, Oita University, Japan, 266 strains from Iran, collected by Abbas Yadegar, and 133 genomes from different parts of the world, including 89 from the Swedish Kalixanda cohort ^53^. Lastly, 1110 worldwide DNA samples were contributed by Professor Mark Achtman. These sequences have also been deposited in Enterobase at the following workspace https://enterobase.warwick.ac.uk/a/108555.

Samples from the Yamaoka lab were sequenced at Novogene Co., Ltd., Beijing, China with the Illumina NOVA PE150 platform and assembled using the SPAdes genome assembler v3.15.3 ^54^ by down sampling read depth to 100 bp, specifying a genome size of 1.6 Mbp and enabling the -- careful option. Of the samples obtained from Prof. Achtman, 920 were sequenced either using the Illumina MiSeq platform at University of Gothenburg or at Novogene Co, Ltd, UK using the Illumina NOVA PE150 platform; and another 190 sequenced at University of Warwick, UK. Remaining sequences were sequenced on the Illumina MiSeq platform at Karolinska Institutet and the University of Gothenburg, Sweden and assembled using the BACTpipe pipeline (https://doi.org/10.5281/zenodo.4742358).

We then filtered out redundant genomes, defined as those sequences with SNP distance <200, and removed low quality genomes based on assembly fragmentation (> 500 contigs), coverage to the 26695 *H. pylori* reference strain (< 70%) and contamination (< 90% *H. pylori*) as predicted by Kraken v2.1.2 ^55^, to obtain a final total of 6870 genomes, correspond to 2663 genomes first reported in this work and 3707 publicly available strains (Table S2).

### Sequence alignment and Core Genome variant calling

Single nucleotide polymorphisms from the core genome (core SNPs) were called using a MUMmer v3.20 ^56^ -based algorithm as previously described ^57^. We first aligned each genome sequence of the entire dataset to the *H. pylori* 26695 reference strain (NC_000915.1) using nucmer v3.1. Then, snp-sites v2.5.2 was used to call all the variants from the whole genome alignment obtained. Variants present in at least 99% of genomes were finally extracted using VCFtools v0.1.17, obtaining a total of 866,840 core SNPs.

### Population assignment

To assign a population to the final dataset, we first defined a reference subset of 285 strains that has been consistently assigned into one of 19 *H. pylori* populations/subpopulations in previous reports ^16,34,58,59^ and that we confirmed by running fineSTRUCTURE ^33^ based on haplotype data from core SNPs, computed for this subset as mentioned above, and using 200,000 iterations of burn-in and Markov chain Monte Carlo (MCMC). Then, this subset was considered as a donor panel to paint each sample of the entire dataset using ChromoPainter v2 ^33^. Genomes were labelled with a population based on the biggest fraction of their genome painted by a population.

### PCA

A PCA (Principal Component Analysis) on the whole dataset was performed using the SNPs extracted from the global alignment file, after linkage disequilibrium-pruning to remove linked SNPs (window size = 50 bp, step size = 10 variants and r2 threshold = 0.1), using the software PLINK (v1.9) ^60^.

### Phylogenetic analysis

To reconstruct the various phylogenetic trees, we used coding sequences that were aligned with strain 26695 (see above). When looking at specific genes (*vacA*, *ureA*, *ureB*), the gene sequences were first obtained from the individual strains annotation file then aligned using MAFFT (v7.505, option --auto) ^61^. The tree from Figure S2A was built using SNPs from all the CDS. The trees from Figure 1B and C (and Figure S2 B and C) were built using SNPs from the undifferentiated (Figure 1B, Figure S2B) and differentiated (Figure 1C, Figure S2C) genes. The trees from Figure S10, S11, S12 and S13 were built using SNPs from specific genes. Additionally, for *ureA* (Fig S12) and *ureB* (Fig S13), the sequences were separated into two types, as some strains had two copies of the gene. The choice of the type of the copy was based on the similarity between the sequences (based on the tree clustering and BLAST results, in particular against *H. cetorum*). Using the various alignments of nucleotide sequences, the Maximum-Likelihood trees were constructed using the FastTree software ^62^ (v2.1.10, option -nt). The trees were then rooted based on a given outgroup that are usually used for *H. pylori*: hpAfrica2 and *H. acinonychis* for the trees looking at the whole genome or at the undifferentiated genes, and the Hardy strains for the trees looking at the differentiated genes. In the case of the trees looking at individual genes, we *H. cetorum* as an outgroup, these genes having been chosen after their sequences were BLASTED against the *H. cetorum* genome (see below). The rooting was done using the R package ape ^63^ (root function).

The population-level trees from Figure 2B were built via a Neighbor-Joining algorithm (R package ape, function nj), using a matrix of the average distance between the populations that were represented in both Hardy and Ubiquitous ecospecies. As an equivalent of *H. acinonychis* (Hardy), we used strains from hpAfrica2. The distances between strains were calculated using the dist.dna function (option model = “raw”) from the ape R ^64^ package.

### FineSTRUCTURE

To further investigate the population structure of Hardy and Ubiquitous strains, we analyzed 295 strains assigned by ChromoPainter to hspIndigenousSAmerica, hspSiberia and hspIndigenousNAmerica *H. pylori* populations by running fineSTRUCTURE with 200,000 iterations of burn-in and Markov chain Monte Carlo (MCMC), using as input the haplotype data prepared with SNPs from the core genome of the 295 strains, considering only those variants present in more than 99% of the samples.

### GWAS, FST and separation into undifferentiated, intermediate, and differentiated genes

Considering the ecospecies as a trait and using the strains from hspSiberia and hspIndigenousNAmerica (244 strains), we performed a GWAS to determine which biallelic core SNPs were significantly associated with the ecospecies. Although Hardy strains were also found in hspIndigenousSAmerica, we choose to remove this population from the GWAS analysis due to the small number (2/49) of Hardy strains. The Ubiquitous strains were coded as 0 (198 strains) while Hardy strains were coded as 1 (46 strains). The GWAS was performed using the R package bugwas (v0.0.0.9000) ^65^, which considers the population structure via a PCA and then uses GEMMA (v0.93) to perform GWAS analysis. Using a standard significance threshold of -log(p) = 5, 4609 out of 285,792 core biallelic SNPs were significantly associated with the Hardy clade.

To reinforce the results obtained by the GWAS, we calculated the per-site FST between the Ubiquitous and Hardy ecospecies via the R package PopGenome ^66^ using the same set of strains and SNPs. We considered a SNP to be differentiated between the Hardy and Ubiquitous ecospecies if it was significantly associated with the ecospecies by the GWAS (-log10(p) > 10) and highly differentiated between the two groups based on its FST value (FST > 0.9). Of the core biallelic SNPs we found 2568 differentiated coding SNPs and 175 differentiated intergenic SNPs. We considered a SNP to be undifferentiated if its -log10(p) was less than 10 and its FST less than 0.5 (265,621 coding and 8950 intergenic SNPs). All the other SNPs (7,756 coding and 591 intergenic) were considered to be intermediate. Following the separation of the SNPs into three classes, we also distinguished three types of genes, based on the genes present in the 26695 genome: differentiated (100 genes, Table S3), containing at least five differentiated SNPs; undifferentiated (1034 genes), with only undifferentiated SNPs; and the remaining genes (443 genes) which we considered as intermediate.

For each strain, we calculated the number of differentiated sites that have the Hardy allele (major allele among the Hardy strains) and compared this number to the genetic distance to the Ubiquitous strains and to the number of Hardy blocks. For a given strain, the distance to the Ubiquitous strains is the average number of differences between the sequences of this strain and sequences of the Ubiquitous strains from hspIndigenousSAmerica, hspSiberia and hspIndigenousNAmerica. The sequences were the sequences aligned on the 26695 sequence and the gaps were removed.

The Hardy blocks were defined based on the differentiated SNPs: for each strain, if two adjacent differentiated SNPs had the same allele and were part of the same gene, we considered that they were part of the same Hardy block, otherwise, they were from different blocks.

### Pangenome analysis

First, we estimated the gene content of each sample with prokka v1.4.6 software ^67^, using the proteome of 26695 as reference. Then, .gff files were used as input for Panaroo’s v1.2.8 ^68^ pangenome pipeline using the strict mode, merging paralogs based on sequence identity of 0.95, length difference of 0.90 and a core threshold of 0.95. For this analysis smaller dataset was used that consisted of all the strains from hspSiberia and hspIndigenousNAmerica (*i.e*., all the Hardy strains were included and all the Ubiquitous strains from the same populations), as well as randomly chosen strains from the other populations (size of the sample dataset: 721 strains).

To detect *cagA*, *vacA*, and *ureAB* homologs in diverse *Helicobacter* species, non-*pylori Helicobacter* genomes were recovered from GenBank or Enterobase (Table S4) and annotated using Prokka. Strains/species were considered if they encoded an intact copy of UreA1B1 (indicating gastric tropism) and available metadata indicated isolation from humans or animals. Metagenome-derived genomes from non-animal hosts were excluded. Host diets were identified using Wikipedia.

Pangenome analysis was performed together with a subset of *H. pylori* strains (Table S2) using panaroo with a 70% sequence identity and 75% sequence coverage cutoff to identify putative homologs. For *vacA*, this was supplemented with additional manual inspection (e.g., incomplete genomes) using Mauve ^69^and Tablet ^70^ and literature information (e.g., *H. cetorum* ^71^).

### Comparison to H. cetorum and phylogenetic analysis of the differentiated genes

We used *H. cetorum* as an outgroup in the study of the differentiated genes. We used the *H. cetorum* strain MIT99-5656 (downloaded the 10/02/2023 from NCBI https://www.ncbi.nlm.nih.gov/data-hub/genome/GCF_000259275.1/). The gene sequences were obtained from the given GenBank file.

A phylogenetic analysis was performed on the differentiated genes, with the *H. cetorum* genome included. First, we obtained the *H. cetorum* gene sequences by BLASTing (blastn v2.11.0) ^72^ a Hardy and Ubiquitous version of each differentiated gene against the *H. cetorum* genome. For the genes that returned at least one hit, the phylogenetic tree of the gene was generated using FastTree, and rooted on the *H. cetorum* sequence (see above).

### Genome structure comparison

Significant structural variations like inversions, gaps, repeats, and gene cluster rearrangement can be easily visualized using a dot plot. To investigate the genome structure similarity and the difference between Hardy and Ubiquitous groups, we used the Gepard program ^73^ (v1.40 jar file from https://cube.univie.ac.at/gepard) to make the dot plot. Different comparisons were considered: Hardy vs. Hardy, Hardy vs. Ubiquitous, and Ubiquitous vs. Ubiquitous, and against *H. cetorum*.

Publicly available genome sequences were downloaded from NCBI GenBank (https://www.ncbi.nlm.nih.gov/genbank/). The Gepard internal DNA substitution matrix (edna.mat) was selected to generate the alignment and plot. The lower color limit was 50 to reduce noise and emphasize significant regions. Window size and word length were default values of 0 and 10, respectively.

### Functional Enrichment analysis

To determine whether particular types of genes were over-represented among the 100 highly differentiated genes, we performed a functional enrichment analysis, using the website DAVID ^74^. Three hypothetical proteins were removed, as they lacked a unique ID. The background set of genes chosen (Table S3 for the list of genes tested and their categories) for the comparison was based on the genes present in *H. pylori* 26695. A term was considered as significant for p < 0.05 after a Benjamini correction for multiple tests.

### dN/dS calculations

For each strain in the dataset, we estimated its dN/dS and dS values to an outgroup population using the Yang and Nielsen method (YN00) of the PAML (v4.9) software ^75,76^. The dN/dS and dS value was averaged over pairwise comparisons with each of the different outgroup strains. The codons that coded for a stop in at least one of the strains were removed from all strains and the dN/dS values were calculated pairwise using the CDS from the sequences aligned to the reference genome (26695; global alignment). Moreover, we separated the values between the undifferentiated and differentiated CDS. We used three different outgroups: hpAfrica2, *H. acinonychis* (which are Hardy strains) and the *H. pylori* Hardy strains, and the outgroup population / ecospecies was not shown in the plots (only the population / ecospecies of the “focal” strain).

### Helicobacter Genomics Consortium

#### Bangladesh

- Hafeza Aftab, Department of Gastroenterology, Dhaka Medical College and Hospital, Dhaka, Bangladesh

#### Bhutan

- Lotay Tshering, Jigme Dorji Wangchuk National Referral Hospital, Thimphu, Bhutan
- Dhakal Guru Prasad, Jigme Dorji Wangchuk National Referral Hospital, Thimphu, Bhutan

#### Congo DR

- Evariste Tshibangu-Kabamba, University of Mbujimayi, Mbujimayi, DR Congo / Osaka Metropolitan University, Osaka, Japan
- Ghislain Disashi Tumba, University of Mbujimayi, Mbujimayi, DR Congo
- Patrick de Jesus Ngoma-Kisoko, University of Kinshasa, Kinshasa, DR Congo
- Antoine Tshimpi-Wola, University of Kinshasa, Kinshasa, DR Congo
- Dieudonné Mumba Ngoyi, University of Kinshasa, Kinshasa, DR Congo / National Institute of Biomedical Research (INRB), DR Congo
- Pascal Tshiamala Kashala, Gastroenterology Service, Astryd Clinics, Kinshasa, DR Congo

#### Dominican Republic

- Modesto Cruz, Instituto de Microbiología y Parasitología (IMPA), Universidad Autónoma de Santo Domingo (UASD), Santo Domingo
- José Jiménez Abreu, Instituto de Microbiología y Parasitología (IMPA), Universidad Autónoma de Santo Domingo (UASD), Santo Domingo
- Celso Hosking, Universidad Autónoma de Santo Domingo, Santo Domingo

#### Finland

- Jukka Ronkainen, Center for Life Course Health Research, University of Oulu; Finland and Primary Health Care Center, Tornio, Finland
- Pertti Aro, Arokero Oy, Tornio, Finland

#### Indonesia

- Titong Sugihartono, Universitas Airlangga, Surabaya, Indonesia
- Ari Fahrial Syam, University of Indonesia, Jakarta, Indonesia
- Langgeng Agung Waskito, Universitas Airlangga, Surabaya, Indonesia
- Hasan Maulahela, University of Indonesia, Jakarta, Indonesia
- Yudith Annisa Ayu Rezkitha, Faculty of Medicine, Muhammadiyah University of Surabaya, Surabaya, Indonesia

#### Iran

- Shaho Negahdar Panirani, Foodborne and Waterborne Diseases Research Center, Research Institute for Gastroenterology and Liver Diseases, Shahid Beheshti University of Medical Sciences, Tehran, Iran.
- Hamid Asadzadeh Aghdaei, Basic and Molecular Epidemiology of Gastrointestinal Disorders Research Center, Research Institute for Gastroenterology and Liver Diseases, Shahid Beheshti University of Medical Sciences, Tehran, Iran.
- Mohammad Reza Zali, Gastroenterology and Liver Diseases Research Center, Research Institute for Gastroenterology and Liver Diseases, Shahid Beheshti University of Medical Sciences, Tehran, Iran
- Nasrin Mirzaei, Foodborne and Waterborne Diseases Research Center, Research Institute for Gastroenterology and Liver Diseases, Shahid Beheshti University of Medical Sciences, Tehran, Iran.
- Saeid Latifi-Navid, Department of Biology, Faculty of Sciences, University of Mohaghegh Ardabili, Ardabil, 5619911367 Iran

#### Japan

- Takeshi Matsuhisa, Nippon Medical School, Tokyo, Japan
- Phawinee Subsomwong, Department of Environmental and Preventive Medicine, Oita University Faculty of Medicine, Yufu, Japan. Department of Microbiology and Immunology, Hirosaki University Graduate School of Medicine, Hirosaki, Japan.
- Hideo Terao, Oita University, Oita, Japan
- Batsaikhan Saruuljavkhlan, Department of Environmental and Preventive Medicine, Oita University Faculty of Medicine, Yufu, Japan.
- Tadashi Shimoyama, Aomori General Health Examination Center, Aomori, Japan
- Nagisa Kinjo, Ryusei Hospital, Naha, Japan
- Fukunori Kinjo, Center for Gastroenterology, Urasoe General Hospital, Urasoe, Japan
- Kazunari Murakami, Department of Gastroenterology, Oita University Faculty of Medicine, Yufu, Japan

#### Myanmar

- Thein Myint, Department of Gastroenterology, Yangon General Hospital, University of Medicine, Yangon, Myanmar
- Than Than Aye, Department of Gastroenterology, Thingangyun Sanpya General Hospital, University of Medicine (2), Thingangyun, Myanmar
- New Ni, Department of Gastroenterology, Mandalay General Hospital, University of Medicine, Mandalay, Myanmar.
- Than Than Yee, Department of GI and HBP Surgery, No. (1) Defense Service General Hospital (1000 Bedded), Mingaladon, Yangon 11021, Myanmar
- Kyaw Htet, Defence Services General Hospital, Yangon, Myanmar

#### Nepal

- Pradeep Krishna Shrestha, Tribhuvan University Teaching Hospital
- Rabi Prakash Sharma, Gastroenterology Department, Maharajgunj Medical Campus, Tribhuvan University Teaching Hospital, Kathmandu, Nepal

#### Sri Lanka

- Jeewantha Rathnayake, Department of Surgery, Teaching Hospital Peradeniya, University of Peradeniya, Kandy 20404, Sri Lanka
- Meegahalande Durage Lamawansa, Department of Surgery, Teaching Hospital Peradeniya, University of Peradeniya, Kandy 20404, Sri Lanka

#### Sweden

- Emilio Rudbeck, Clinical Genomics Gothenburg, Bioinformatics and Data Centre, University of Gothenburg, Sweden
- Lars Agreus, Division of Family Medicine, Karolinska Institutet, Stockholm, Sweden
- Anna Andreasson, Stress Research Institute, Department of Psychology, Faculty of Social Sciences, Stockholm University, Sweden
- Lars Engstrand, Center for Translational Microbiome Research, Department for Microbiology, Tumor, and Cell Biology, Karolinska Institutet, Stockholm, Sweden

#### Thailand

- Varocha Mahachai, GI and Liver Center, Bangkok Medical Center, Bangkok, Thailand
- Thawee Ratanachu-Ek, Department of Surgery, Rajavithi Hospital, Bangkok, Thailand
- Kammal Kumar Pawa, Department of Medicine, Chulabhorn International College of Medicine (CICM) at Thammasat University, Pathumthani, Thailand

#### Vietnam

- Tran Thi Huyen Trang, 108 Military Central Hospital, Hanoi, Vietnam
- Tran Thanh Binh, Tam Anh General Hospital, Ho Chi Minh city, Vietnam
- Ho Dang Quy Dung, Cho Ray Hospital, Ho Chi Minh, Vietnam
- Vu Van Khien, 108 Military Central Hospital, Hanoi, Vietnam
- Dou Narith, Department of Endoscopy, Cho Ray Phnom Penh Hospital, Phnom Penh 12357, Cambodia.

## Ethics & Inclusion statement

The Helicobacter genomics consortium includes gastroenterologists and researchers from several developing countries. Its aim is to characterize genetic diversity of Helicobacter in human populations across the world, and correlations with gastric disease. In Bangladesh, Bhutan, Congo DR, Dominican Republic, Indonesia, Japan, Myanmar, Nepal, Sri Lanka, Thailand, and Vietnam, all preparation for endoscopy survey was performed by local researchers (persons in the consortium and PhD students at Oita University). Ethical permission for the collection of human gastric biopsy material had been obtained for all cohorts, including informed consent from the participating individuals. Endoscopy was performed by local physicians and Dr Yamaoka. The culture of the bacteria, DNA extraction, next-generation sequencing, and basic genetic analysis in these countries were performed at Oita University, principally by doctoral students who were locally recruited in several of the countries and are coauthors of the paper. In addition, the doctors and scientists involved in this consortium are actively involved in the research process and kept up to date with its findings. This training and dissemination effort will help to spread both genomics knowledge and best practice for treating gastric illness from Japan, where there has been considerable success in mitigating the burden of gastric cancer and other conditions, to other less developed nations.

The study protocol (Iranian strains) was approved by the Institutional Ethical Review Committee of the Research Institute for Gastroenterology and Liver Diseases at Shahid Beheshti University of Medical Sciences, Tehran, Iran (IR.SBMU.RIGLD.REC.1395.878). All experiments were performed in accordance with relevant guidelines and regulations recommended by the institution. Written informed consents were obtained from all enrolled subjects and/or their legal guardians prior to sample collection.

The sampling of the DNA used to generate the new Siberian genomes has been detailed previously ^16^. The remaining new genomes were also from previously collected *H. pylori* DNA, e.g. ^34,35,59^. The study protocol for the Swedish Kalixanda genomes was approved by Umeå University ethics committee, and the study was conducted in accordance with the Helsinki declaration.

Our finding of a highly distinct Hardy ecospecies, with genetic differences at many known virulence factors has potential implications for treatment of infected individuals in many Indigenous communities, which are known to have a high gastric disease burden. However, the pathogenicity profile, either in single or mixed infection is currently unknown. We are maintaining and developing contacts with researchers working in communities where Hardy strains have been isolated from, with a view to consulting with these communities as soon as clinically relevant information becomes available, as well as to performing future clinical studies.

## Data Availability Statement

The list of strains used are provided in the Supplementary Tables S1 and S2. The newly sequenced strains have been deposited on Enterobase and will be made available upon publication. The public strains can be found on Enterobase, NCBI, BIGs and figshare.

The scripts for most of the analysis (except for the fineSTRUCTURE and chromosome painting ones) are available on github (https://github.com/EliseTourrette/HpEcospecies_Tourrette2023). In particular, these scripts contain the detail on how most of the analysis was done and the parameters chosen.

## Acknowledgements

We thank Mark Achtman for providing strain DNA and Jay Solnick for information on primate strains.

## Supplementary Figure Legends

Figure S1:

**Origin of the different strains of our global dataset.** The size of the pie charts shows the number of isolates from the country, with areas scaling logarithmically with sample size. The pie charts show the proportion of isolates assigned to each *H. pylori* population.

Figure S2:

**Phylogenetic trees for all strains in the dataset.** The branches are colored based on their population. Hardy strains are represented with a circle while the other strains from the same populations are represented with a cross. The primate strains are represented with squares. Phylogenetic trees for all the genes (A). Some strains from hspIndigenousNAmerica, hspIndigenousSAmerica and hspSiberia do not cluster with their expected population. For undifferentiated genes (B) and for the differentiated genes (C). The branches are colored based on the population and the strains from the Hardy clade are indicated with a dot.

Figure S3:

**First two components of the Principal Components Analysis (PCA) from the entire dataset.** Strains are colored based on their population and the strains from the Hardy ecospecies are represented by a dot while the crosses represent Ubiquitous strains. Squares and circles respectively indicate primate and *H. acinonychis* strains.

Figure S4:

**FineSTRUCTURE analysis of the strains from hspSiberia, hspIndigenousNAmerica and hspIndigenousSAmerica.** The strains from the Hardy clade are highlighted by red shading, overlaying the dendrogram on the left of the plot, while the Ubiquitous strains are highlighted with blue. FineSTRUCTURE uses an *in silico* chromosome painting algorithm to fit each strain as a mosaic of nearest neighbors, chosen from the other strains in the dataset. Each row shows the coancestry vector for one strain, which is a count of the number of segments of DNA used in the painting from each of the other strains in the dataset. High coancestry between strains implies that they are nearest neighbors for many segments of the genome and hence share genetic material from a common gene pool. FineSTRUCTURE based clustering is more sensitive to recent gene flow than clustering using genetic distances.

Figure S5:

**Manhattan plot resulting from a GWAS analysis of the Hardy vs Ubiquitous strains from hspSiberia and hspIndigenousNAmerica.** The green line represents the significance threshold (- log10(p) = 10). Points are coloured based on FST (fixation index between Hardy and Ubiquitous ecospecies) values (red: FST > 0.9, blue: FST 0.5 - 0.9, grey: FST < 0.5). Half points at the top of the plot indicate an estimated p-value of zero and FST of one.

Figure S6:

**Average ancestry profiles of Global *H. pylori*.** (A) With hpSahul donors and (B) without hpSahul donors. Close-up of Hardy primates and hpSahul strains (C) with hpSahul donors and (D) without hpSahul donors. Although the primate strains do not correspond to any hpSahul strains present in the data (based on their ancestry profile in the presence of an hpSahul donor), they can still be assigned to hpSahul (based on their ancestry profile without an hpSahul donor).

Figure S7:

**Dot plot comparisons between genomes within and between ecospecies**. The genomes of two hspIndigenousNAmerica, one Hardy and one Ubiquitous strain were plotted against the genome of strains more or less distantly related, from left to right: hspIndigenousNAmerica, hspIndigenousSAmerica, hspSiberia, hpAsia2, hpEurope, hpAfrica1, hpAfrica2, *H. acinonychis* and *H. cetorum;* and from top to bottom: Hardy vs Hardy strains, Hardy vs Ubiquitous strains and Ubiquitous vs Ubiquitous strains. Comparison between identical genomes would give single diagonal line, with breaks indicating rearrangements and differences in genome content. The presence of several small lines indicates that there are many rearrangements between the two genomes being compared. On the contrary, comparisons with long lines means highly similar genomes. For more details on how the comparisons were made, see the paragraph "Genome structure comparison" in the Method section.

Figure S8:

**Pairwise dN/dS values to relevant outgroups**. dN/dS vs dS (A,B) dN/dS vs dS between the different populations (hpAfrica2 excluded) and hpAfrica2 for the undifferentiated (A) and differentiated (B) genes. Thus, all comparisons involve hpAfrica2 strains and the dots are colored based on the non-hpAfrica2 population. In addition, the shape represents the ecospecies of the non-hpAfrica2 strain, the dots represent Hardy strains while the crosses represent the Ubiquitous strains; the primates and *H. acinonychis* strains are indicated with squares and circles, respectively. (C,D) dN/dS vs dS between the Ubiquitous and Hardy strains for the undifferentiated (C) and differentiated (D) genes (subplots based on whether the Hardy strains were *H. acinonychis* or non-*H. acinonychis*). For the C and D subplots, all comparisons involve one Hardy (*H. acinonychis* or non-*H. acinonychis*) and one Ubiquitous strain, and the dots are colored based on the Ubiquitous strain population. In all cases, each dot represents the value for a non-outgroup strain, averaged over their values when compared against the different outgroup strains (the outgroups are hpAfrica2 for subplots A and B and *H. acinonychis* or Hardy *H. pylori* for subplots C and D).

Figure S9:

**Number of Hardy alleles per strain against the number of Hardy blocks.** The dots represent the Hardy strains, and the points are colored based on their population. The outliers from Fig. 3B are shown with a x (the inset zoom around these outliers). The primate and *H. acinonychis* strains are indicated by squares and circles, respectively.

Figure S10: **Phylogenetic trees of the sequences of differentiated genes that returned at least one hit when blasted against the *H. cetorum* genome.** The trees can be separated into four main scenarios: (A) The two ecospecies diverged after the divergence between *H. cetorum* and *H. pylori*, but before the hpAfrica2 branch split (scenario 1). In this scenario, *H. cetorum* is the deepest branch in the tree with the next deepest branch separating Hardy and Ubiquitous strains. (B) The divergence takes place after the split with hpAfrica2 (scenario 2). This scenario is similar to (A) except that *H. acinonychis* is the second deepest branch after *H. cetorum*, meaning that human and primate strains cluster together. (C) Presence of gene flow (scenario 3). In this scenario, there are distinct Hardy and Ubiquitous clusters but some Hardy or Ubiquitous strains have the wrong type. (D) Variant of scenario 1 but two copies are present. (E) The polymorphism is older than the divergence with *H. cetorum* (scenario 4). In this case, the deepest branch of the tree is between Hardy and Ubiquitous, with *H. cetorum* clustering with one or other.

Figure S11:

**ML phylogenetic tree of *vacA*, based on nucleotide sequences from complete *H. pylori* genomes and *H. cetorum* (AFI05407.1).** The branches are colored based on their population and the Hardy strains are represented with a dot. Contrary to the *vacA* tree that is shown in Figure S10D, this tree was built using only complete genomes (in most of the non-complete genomes, *vacA* appears to be highly fragmented due to assembly issues). Only *one* copy of vacA from *H. cetorum* was included as they all clustered on the same branch. The leaves are labeled with strain name, plus the number of the copy of the gene in that strain (i.e., .1, and if there are more than one copy, .2 and .3). Note that, for the Hardy strains, the copies that are closer to the root (generally one copy per Hardy strain) are more *H. pylori-*like than the copies farther away.

Figure S12:

**Phylogenetic tree of *ureA* sequences for a sample of strains.** (A) For all *ureA* copies, (B) for the first copy, which is conserved in all *H. pylori* strains and (C) for the second copy that is only present in the Hardy strains and in some of the Ubiquitous strains from the same populations as the Hardy ones. The strains are colored based on their population, a dot at the end of the branch indicates a Hardy strain. *H. cetorum* sequences are also included.

Figure S13:

**Phylogenetic tree of *ureB* sequences for a sample of strains.** (A) For all *ureB* copies, (B) for the first copy that is present in all strains and (C) for the second copy that is only present in the Hardy strains and in some of the Ubiquitous strains from the same populations as the Hardy ones. The strains are colored based on their population, a dot at the end of the branch indicates a Hardy strain. *H. cetorum* sequences are included.

## Supplementary Tables

Supplementary Table 1

**Data availability**

Supplementary Table 2

**Genomes used in the study**

Supplementary Table 3

**List of genes that are differentiated, undifferentiated and intermediate between Hardy and Ubiquitous ecospecies**

Supplementary Table 4

**List of animal *Helicobacter* strains.**

Supplementary Table 5

**List of significantly enriched functions amongst the 100 differentiated genes**

