## Supplementary Figures 1 to 13 for "An ancient ecospecies of *Helicobacter pylori* found in Indigenous populations and animal adapted lineages"

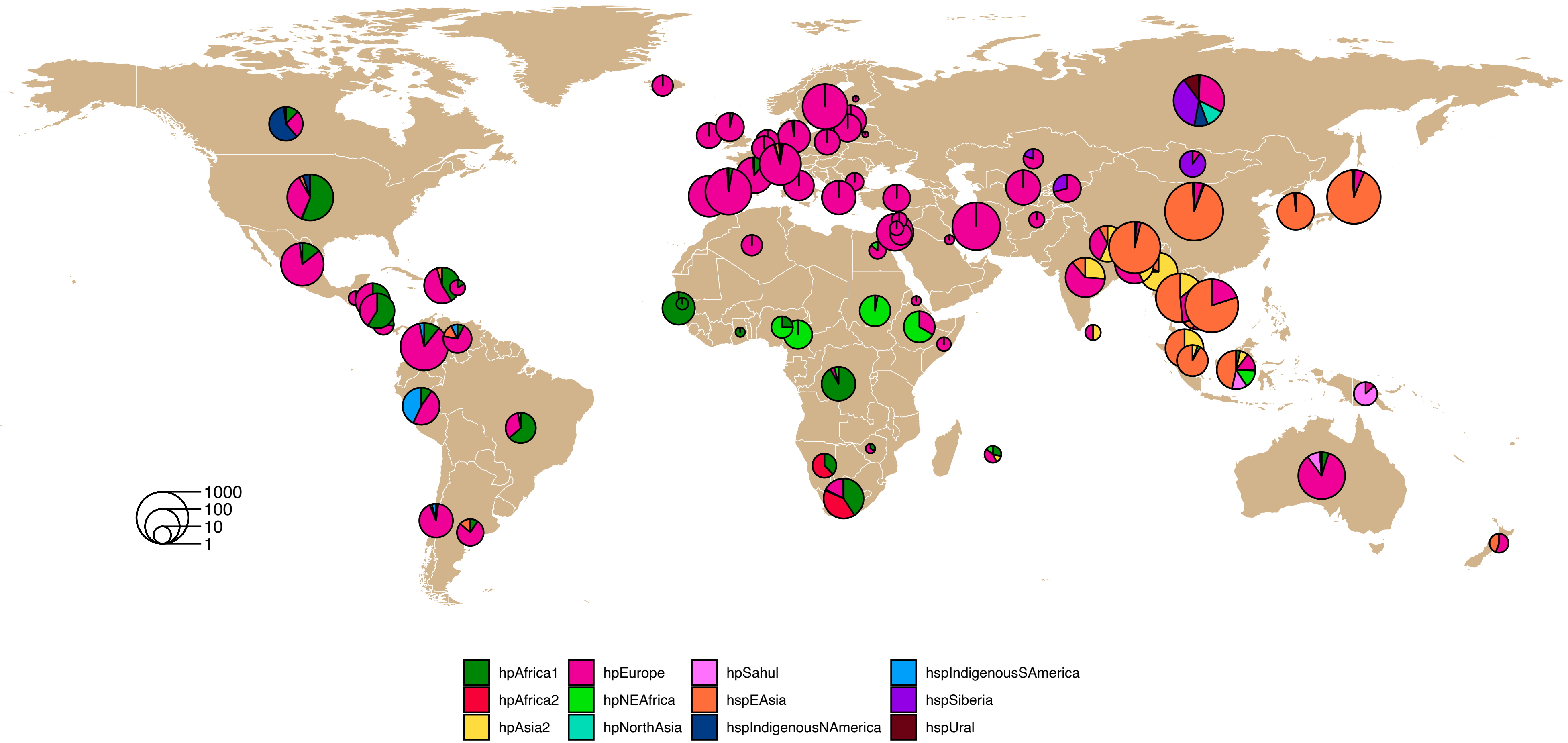


Figure S1:

**Origin of the different strains of our global dataset.** The size of the pie charts shows the number of isolates from the country, with areas scaling logarithmically with sample size. The pie charts show the proportion of isolates assigned to each *H. pylori* population.

A. Tree for the whole genome, rooted by *H. acinonychis*

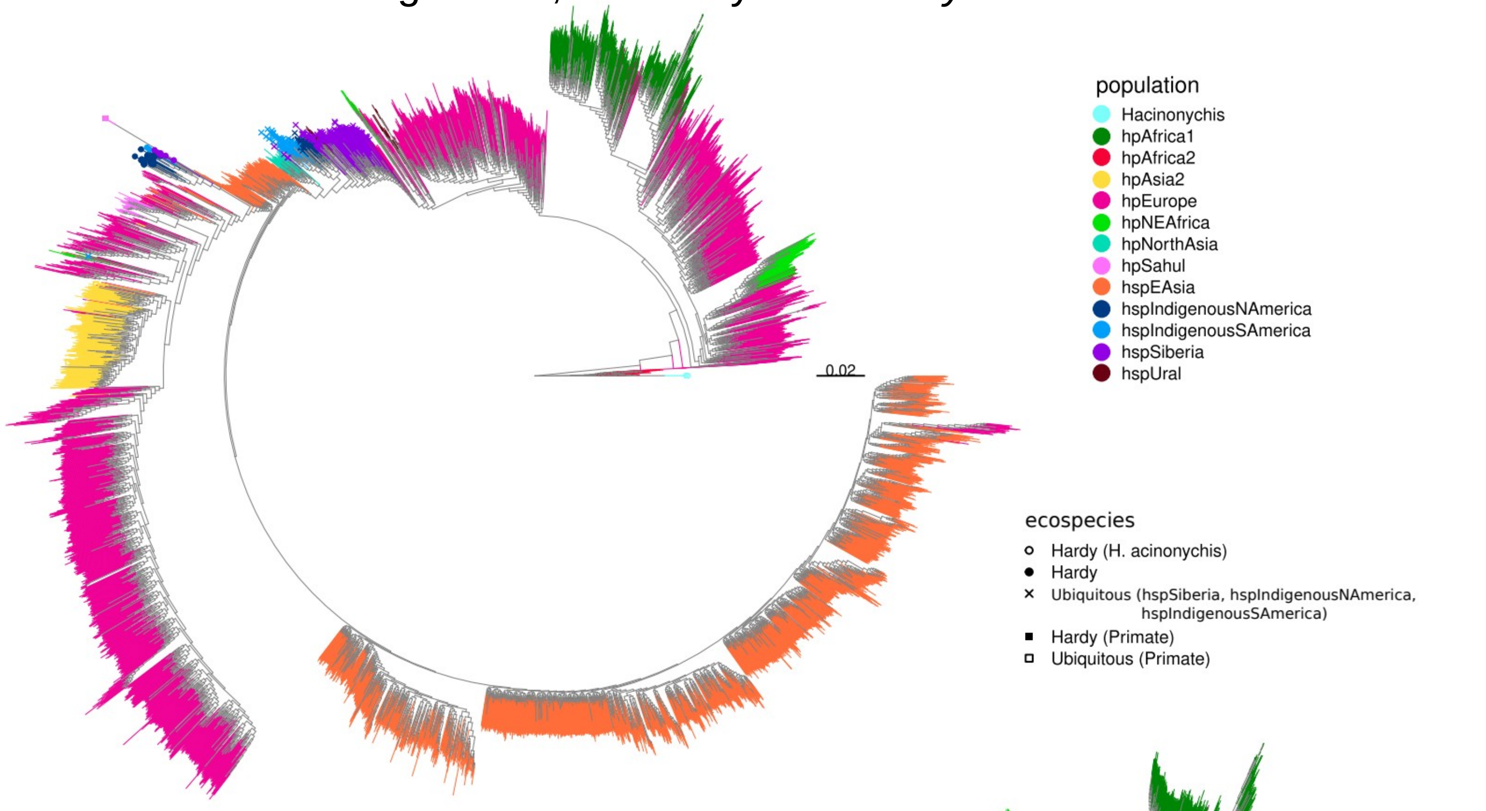


B. Tree for the undifferentiated genes, rooted by *H. acinonychis*

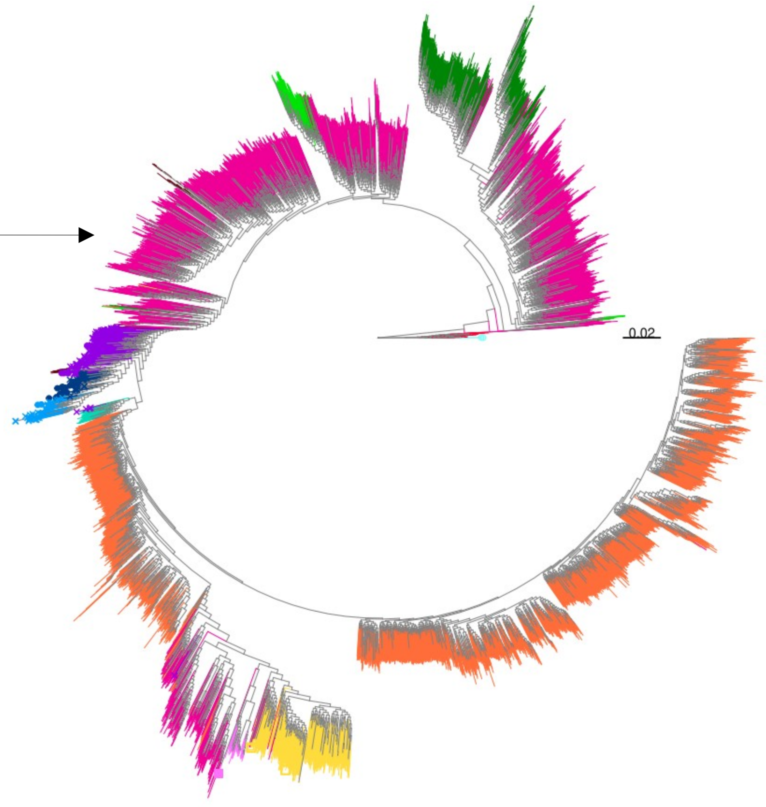


C. Tree for the differentiated genes, rooted by the Hardy strains

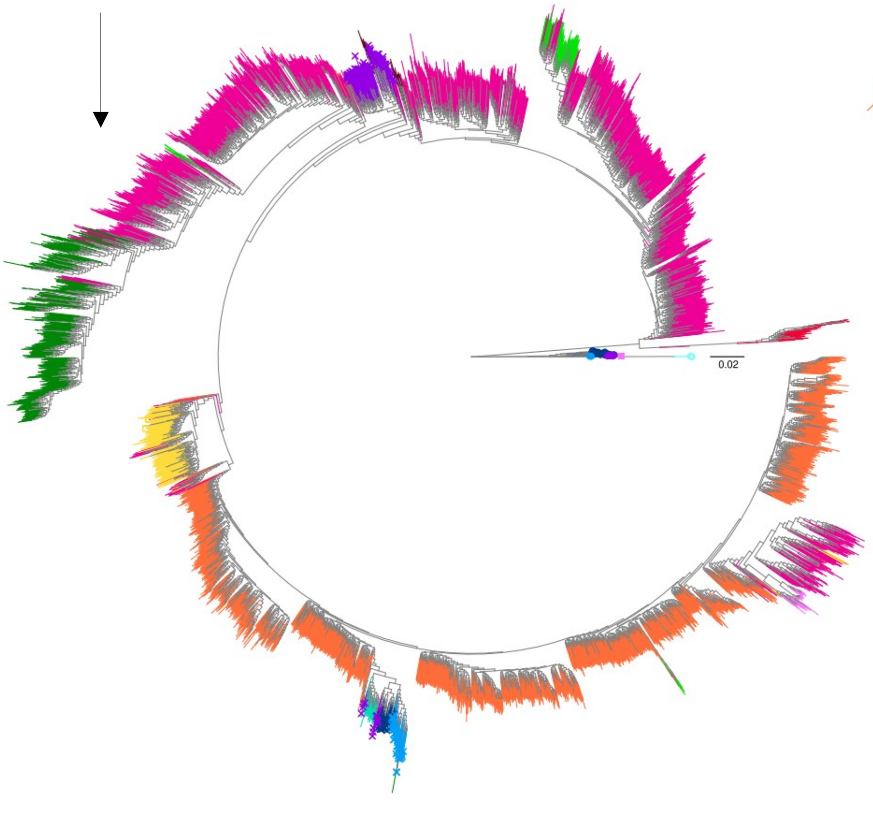


Figure S2:
**Phylogenetic trees for all strains in the dataset.** The branches are colored based on their population. Hardy strains are represented with a circle while the other strains from the same populations are represented with a cross. The primate strains are represented with squares.
Phylogenetic trees for all the genes (A). Some strains from hspIndigenousNAmerica, hspIndigenousSAmerica and hspSiberia do not cluster with their expected population. For undifferentiated genes (B) and for the differentiated genes (C). The branches are colored based on the population and the strains from the Hardy clade are indicated with a dot.

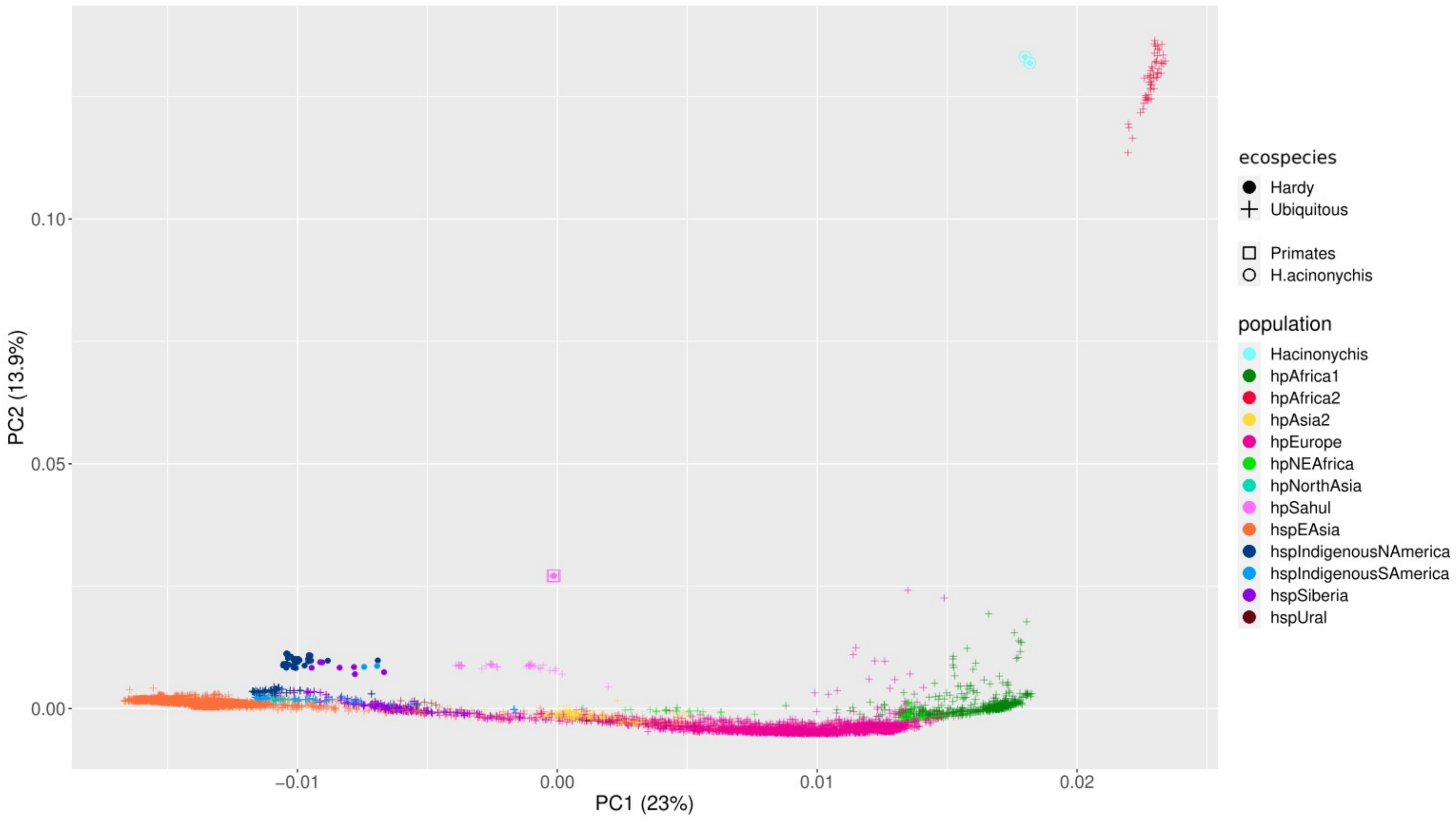


Figure S3:

**First two components of the Principal Components Analysis (PCA) from the entire dataset.** Strains are colored based on their population and the strains from the Hardy ecospecies are represented by a dot while the crosses represent Ubiquitous strains. Squares and circles respectively indicate primate and *H. acinonychis* strains.

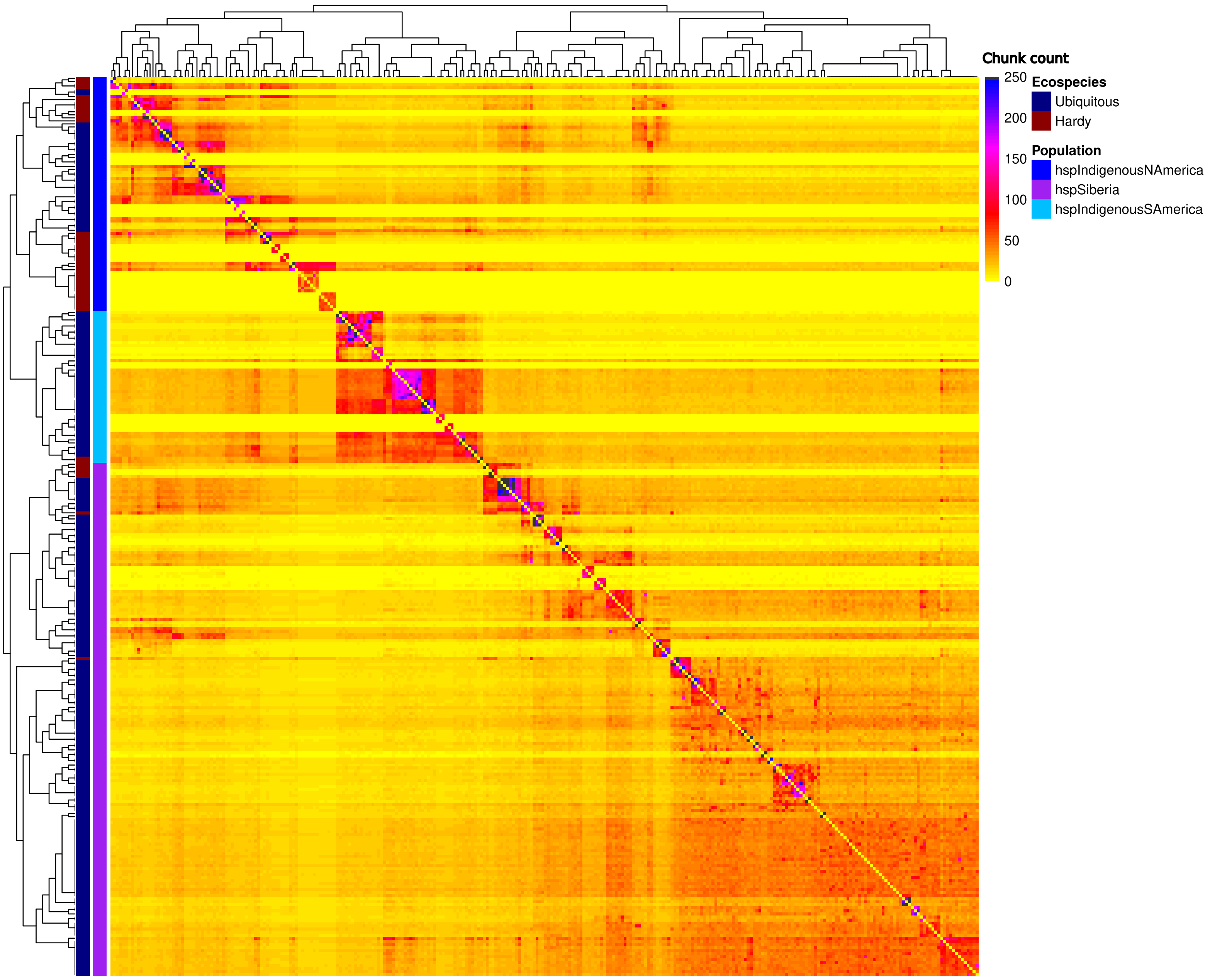


Figure S4:
**FineSTRUCTURE analysis of the strains from hspSiberia, hspIndigenousNAmerica and hspIndigenousSAmerica.** The strains from the Hardy clade are highlighted by red shading, overlaying the dendrogram on the left of the plot, while the Ubiquitous strains are highlighted with blue. FineSTRUCTURE uses an *in silico* chromosome painting algorithm to fit each strain as a mosaic of nearest neighbors, chosen from the other strains in the dataset. Each row shows the coancestry vector for one strain, which is a count of the number of segments of DNA used in the painting from each of the other strains in the dataset. High coancestry between strains implies that they are nearest neighbors for many segments of the genome and hence share genetic material from a common gene pool. FineSTRUCTURE based clustering is more sensitive to recent gene flow than clustering using genetic distances.

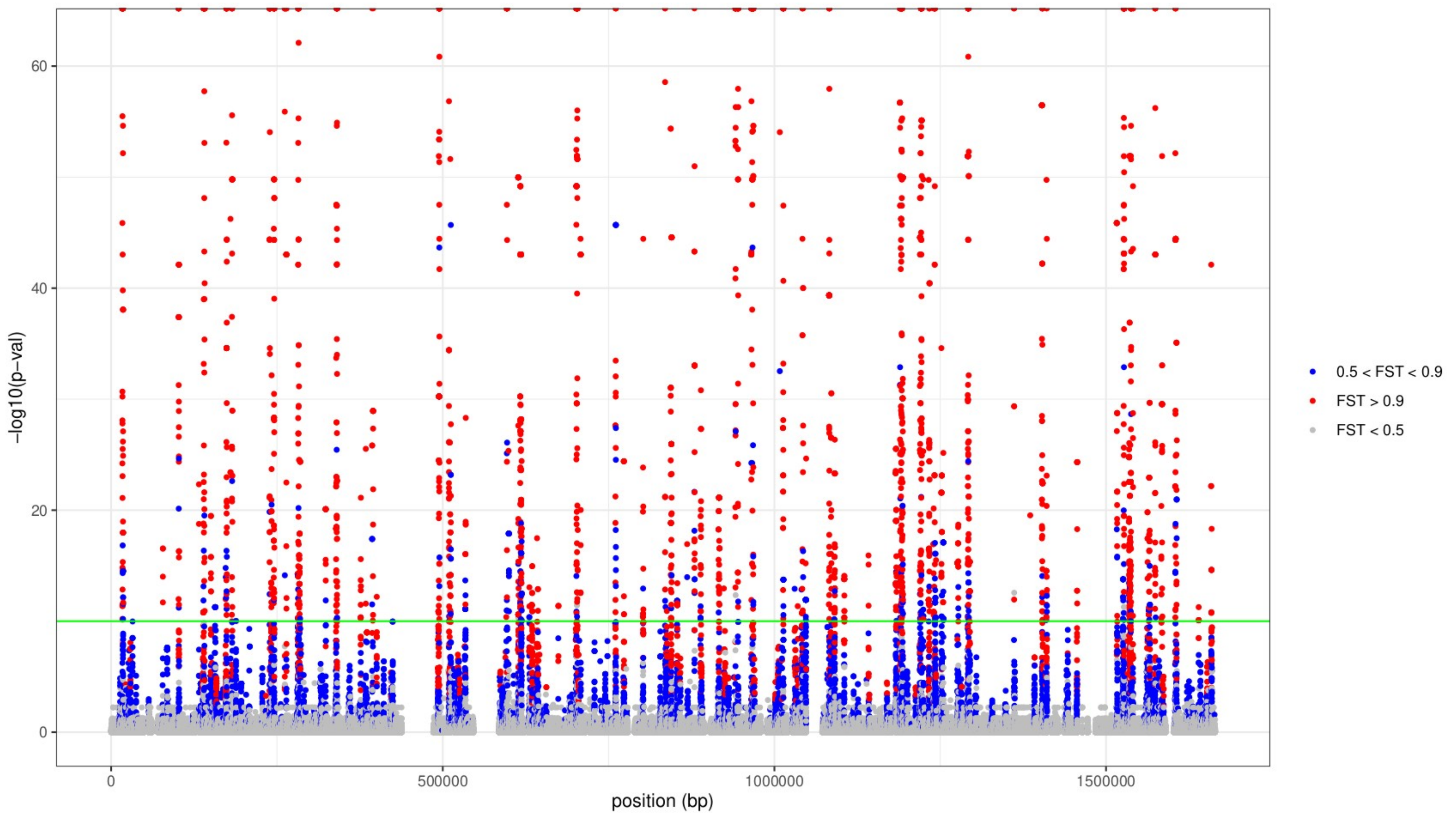


Figure S5:

**Manhattan plot resulting from a GWAS analysis of the Hardy vs Ubiquitous strains from hspSiberia and hspIndigenousNAmerica.** The green line represents the significance threshold (-log10(p) = 10). Points are colored based on their FST (fixation index between Hardy and Ubiquitous ecospecies) values (red: FST > 0.9, blue: FST 0.5 - 0.9, grey: FST < 0.5). Half points at the top of the plot indicate an estimated p-value of zero and FST of one.

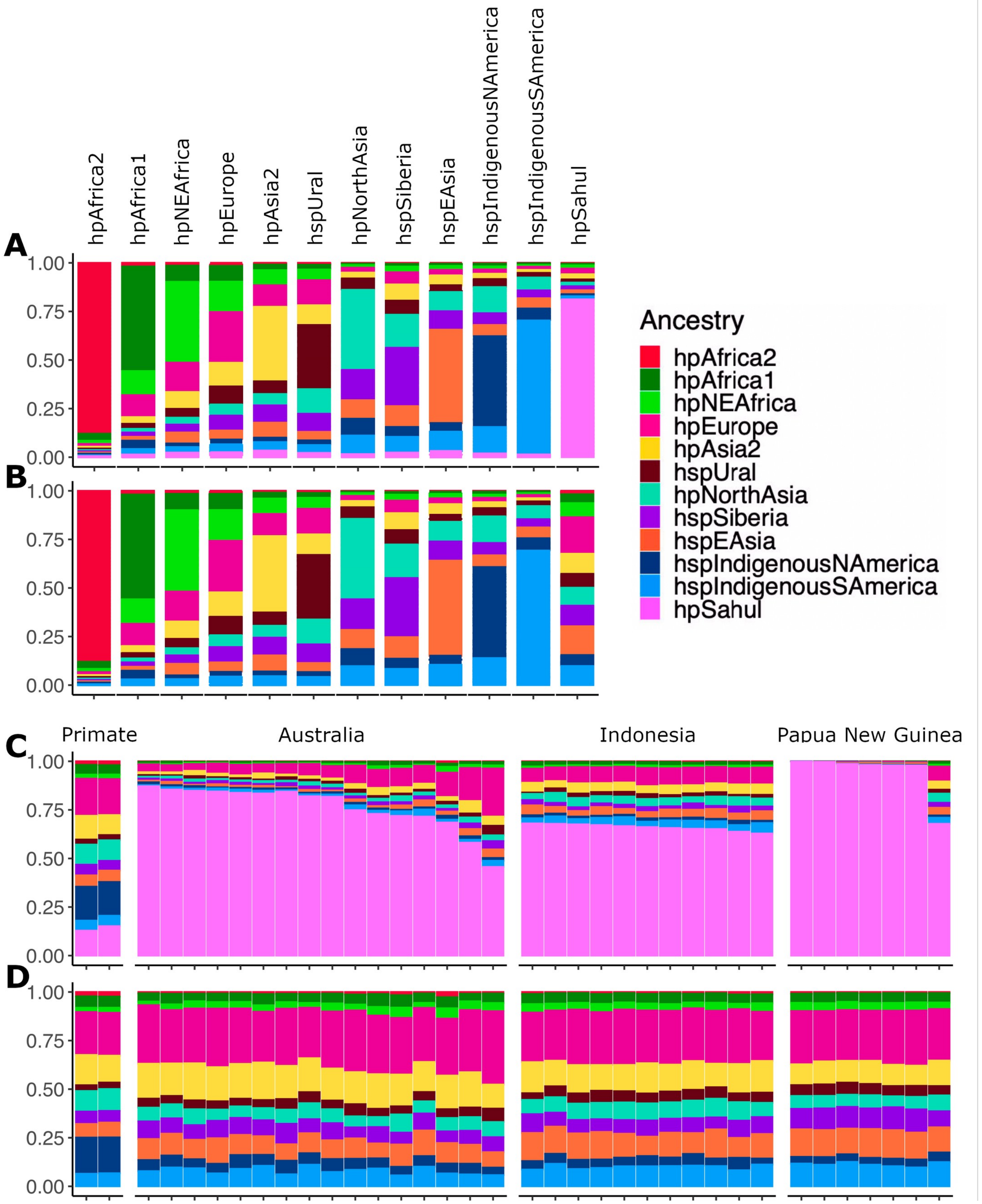


Figure S6:

**Average ancestry profiles of Global *H. pylori.*** (A) With hpSahul donors and (B) without hpSahul donors. Close-up of Hardy primates and hpSahul strains (C) with hpSahul donors and (D) without hpSahul donors. Although the primate strains do not correspond to any hpSahul strains present in the data (based on their ancestry profile in the presence of an hpSahul donor), they can still be assigned to hpSahul (based on their ancestry profile without an hpSahul donor).

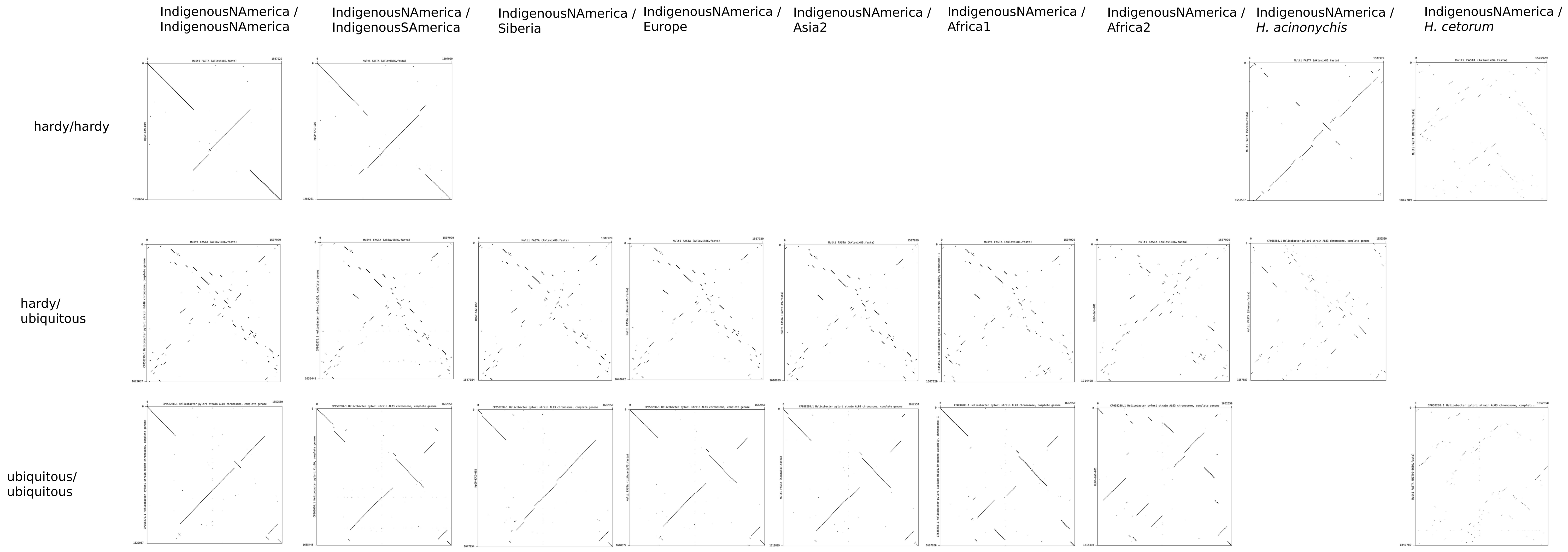


Figure S7:

**Dot plot comparisons between genomes within and between ecospecies**. The genomes of two hspIndigenousNAmerica, one Hardy and one Ubiquitous strain were plotted against the genome of strains more or less distantly related, from left to right: hspIndigenousNAmerica, hspIndigenousSAmerica, hspSiberia, hpAsia2, hpEurope, hpAfrica1, hpAfrica2, *H. acinonychis* and *H. cetorum;* and from top to bottom: Hardy vs Hardy strains, Hardy vs Ubiquitous strains and Ubiquitous vs Ubiquitous strains. Comparison between identical genomes would give single diagonal line, with breaks indicating rearrangements and differences in genome content. The presence of several small lines indicates that there are many rearrangements between the two genomes being compared. On the contrary, comparisons with long lines means highly similar genomes. For more details on how the comparisons were made, see the paragraph "Genome structure comparison" in the Method section.

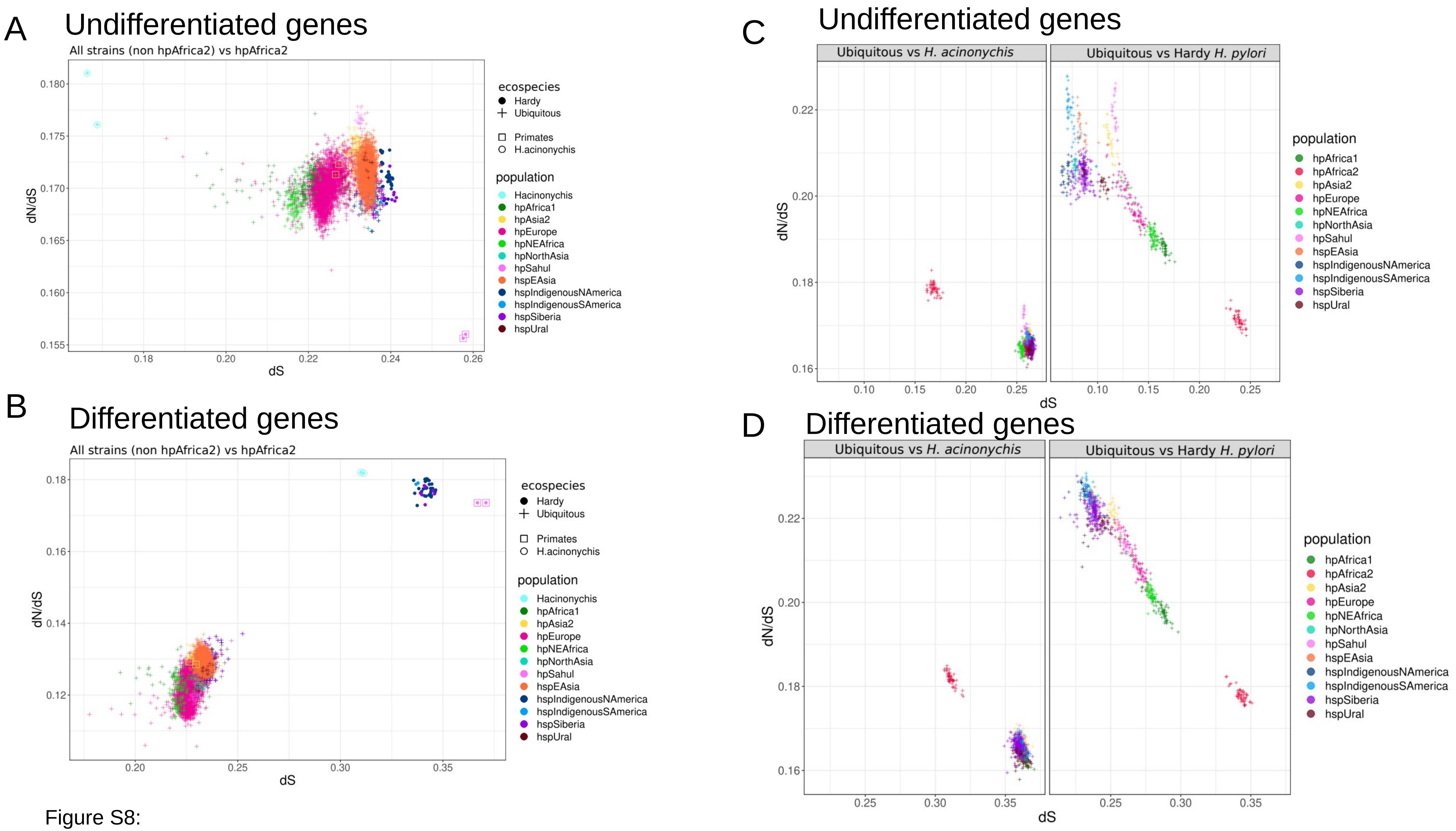


Figure S8:

**Pairwise dN/dS values to relevant outgroups.** dN/dS vs dS (A,B) dN/dS vs dS between the different populations (hpAfrica2 excluded) and hpAfrica2 for the undifferentiated (A) and differentiated (B) genes. Thus, all comparisons involve hpAfrica2 strains and the dots are colored based on the non-hpAfrica2 population. In addition, the shape represents the ecospecies of the non-hpAfrica2 strain, the dots represent Hardy strains while the crosses represent the Ubiquitous strains; the primates and *H. acinonychis* strains are indicated with squares and circles, respectively. (C,D) dN/dS vs dS between the Ubiquitous and Hardy strains for the undifferentiated (C) and differentiated (D) genes (subplots based on whether the Hardy strains were *H. acinonychis* or non-*H. acinonychis*). For the C and D subplots, all comparisons involve one Hardy (*H. acinonychis* or non-*H. acinonychis*) and one Ubiquitous strain, and the dots are colored based on the Ubiquitous strain population. In all cases, each dot represent the value for a non-outgroup strain, averaged over their values when compared against the different outgroup strains (the outgroups are hpAfrica2 for subplots A and B and *H. acinonychis* or Hardy *H. pylori* for subplots C and D).

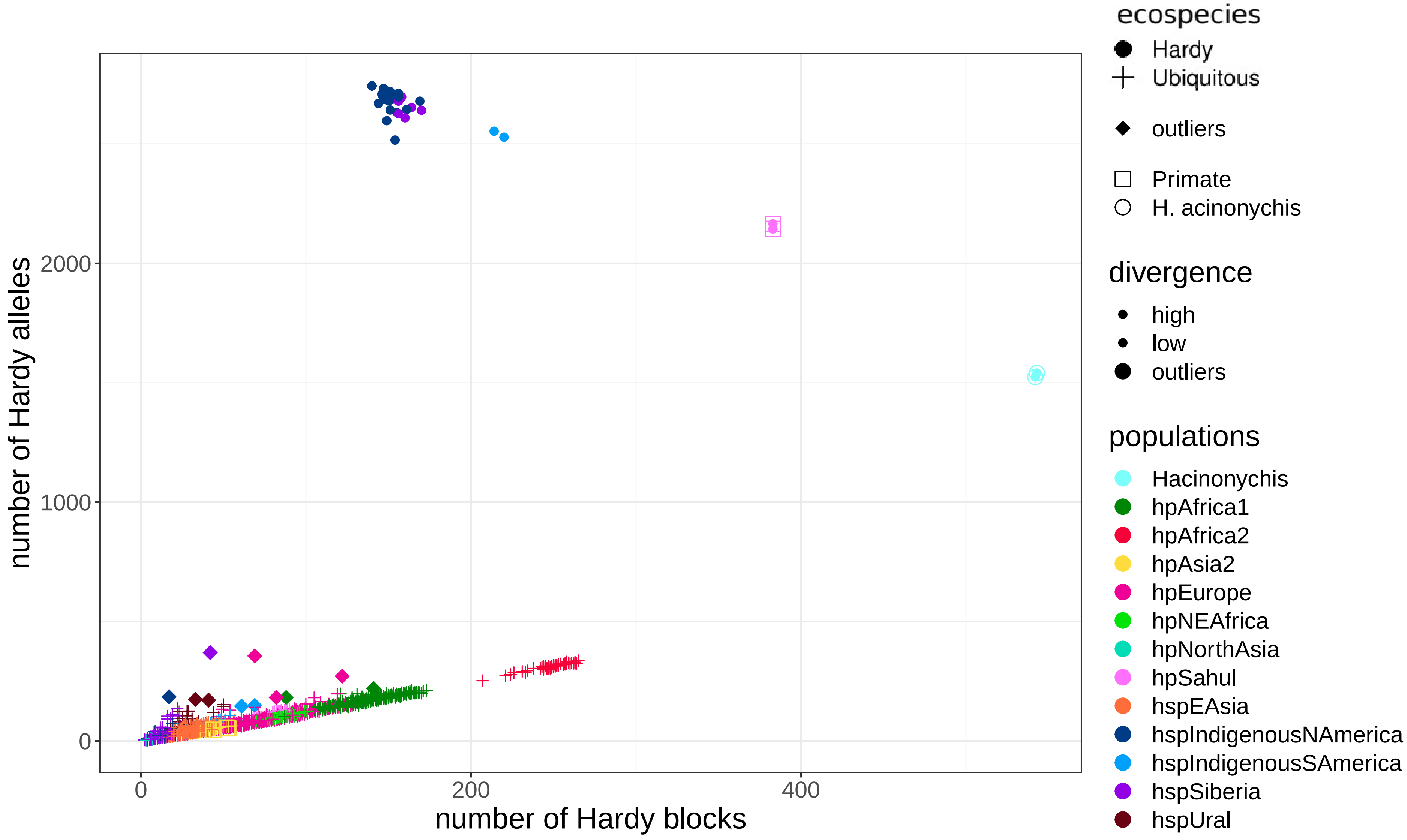


Figure S9:
**Number of Hardy alleles per strain against the number of Hardy blocks.** The dots represent the Hardy strains, and the points are colored based on their population. The outliers from Fig. 3B are shown with a diamonds. The primate and *H. acinonychis* strains are indicated by squares and circles, respectively.

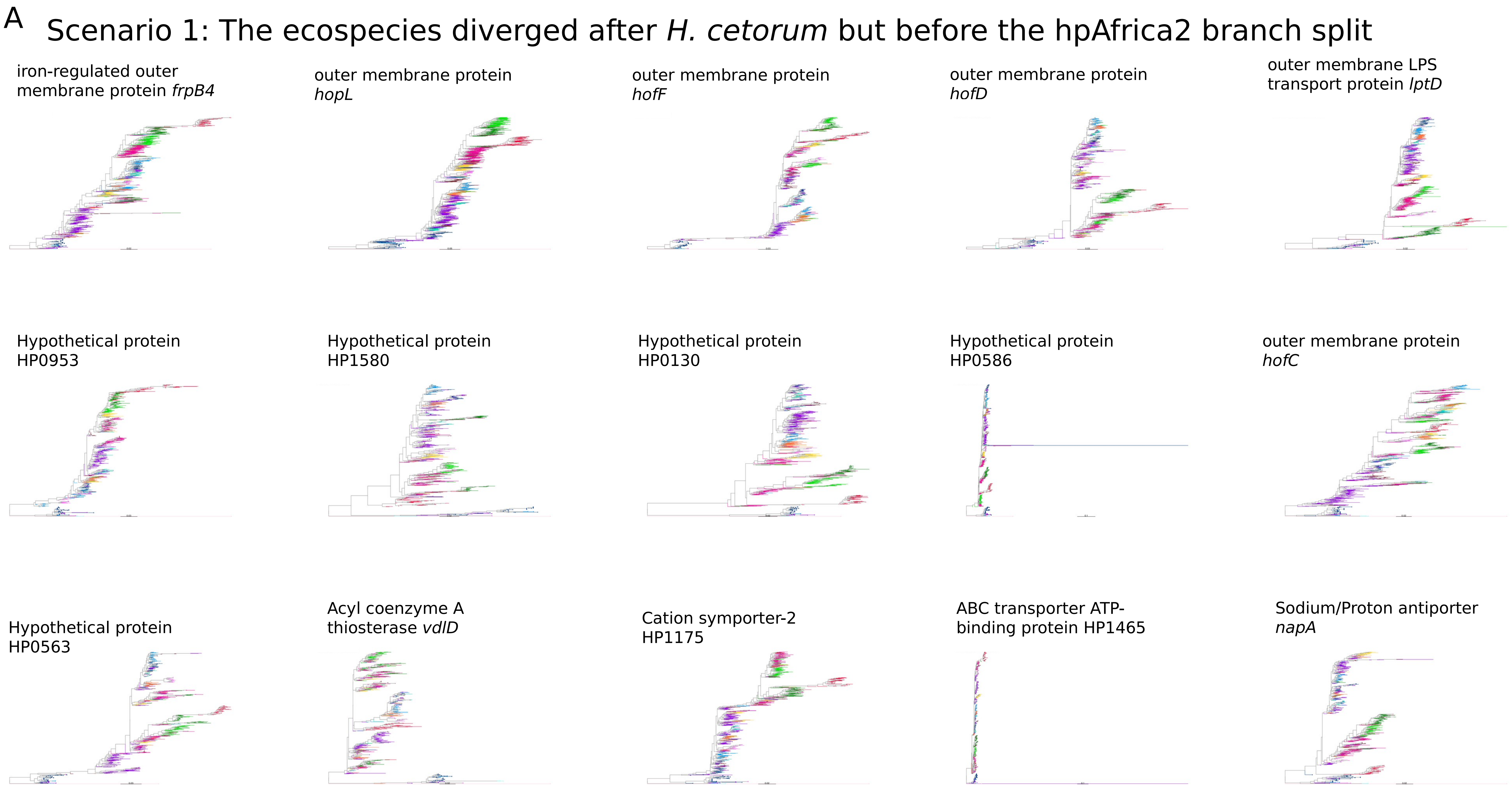
A
Scenario 1: The ecospecies diverged after *H. cetorum* but before the hpAfrica2 branch split
iron-regulated outer membrane protein *frpB4*
outer membrane protein *hopL*
outer membrane protein *hofF*
outer membrane protein *hofD*
outer membrane LPS transport protein *lptD*
Hypothetical protein HP0953
Hypothetical protein HP1580
Hypothetical protein HP0130
Hypothetical protein HP0586
outer membrane protein *hofC*
Hypothetical protein HP0563
Acyl coenzyme A thiosterase *vdlD*
Cation symporter-2 HP1175
ABC transporter ATP-binding protein HP1465
Sodium/Proton antiporter *napA*

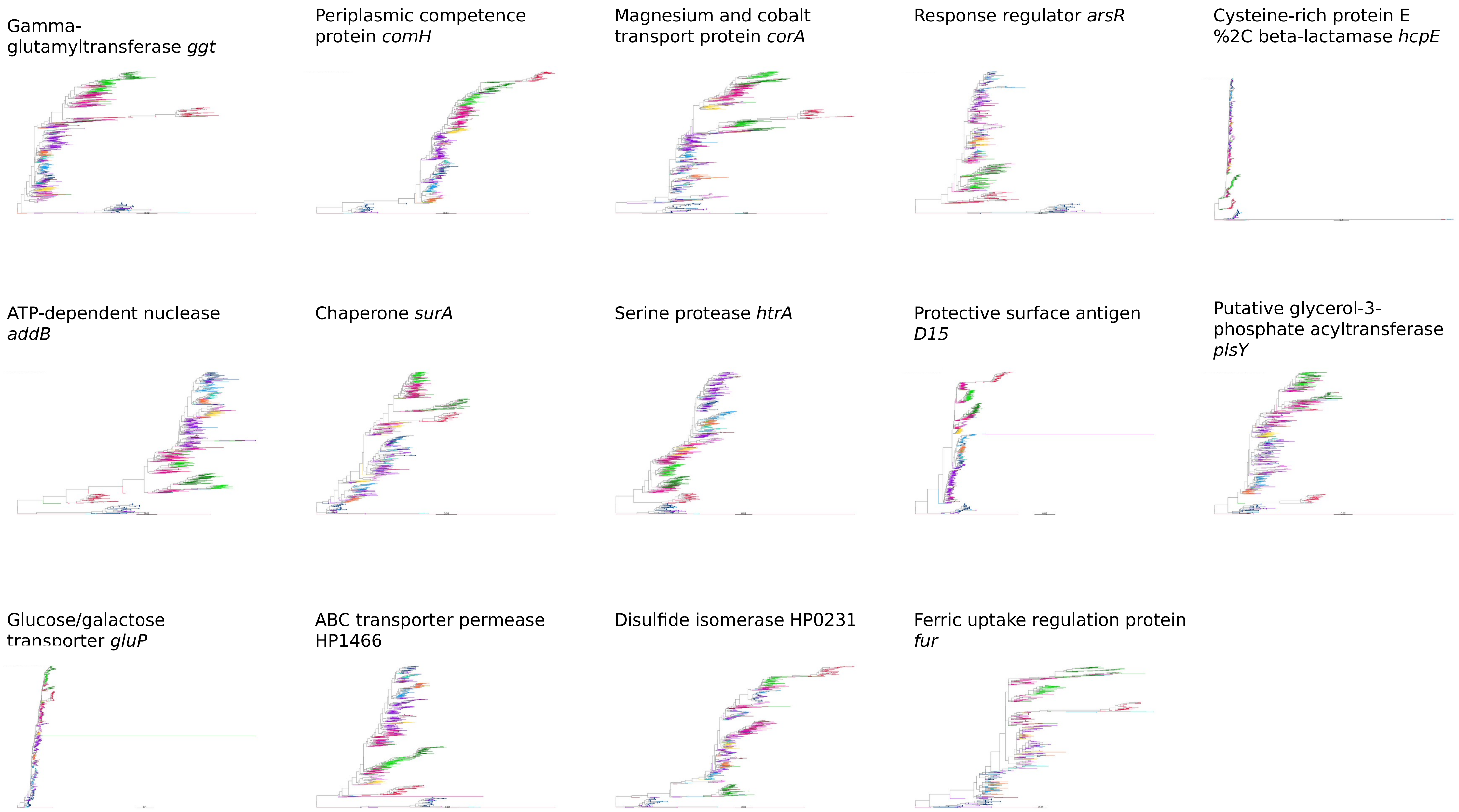
Gamma-glutamyltransferase *ggt*
Periplasmic competence protein *comH*
Magnesium and cobalt transport protein *corA*
Response regulator *arsR*
Cysteine-rich protein E %2C beta-lactamase *hcpE*
ATP-dependent nuclease *addB*
Chaperone *surA*
Serine protease *htrA*
Protective surface antigen *D15*
Putative glycerol-3-phosphate acyltransferase *plsY*
Glucose/galactose transporter *gluP*
ABC transporter permease HP1466
Disulfide isomerase HP0231
Ferric uptake regulation protein *fur*

B

#### Scenario 2: The divergence takes place after the divergence with hpAfrica2

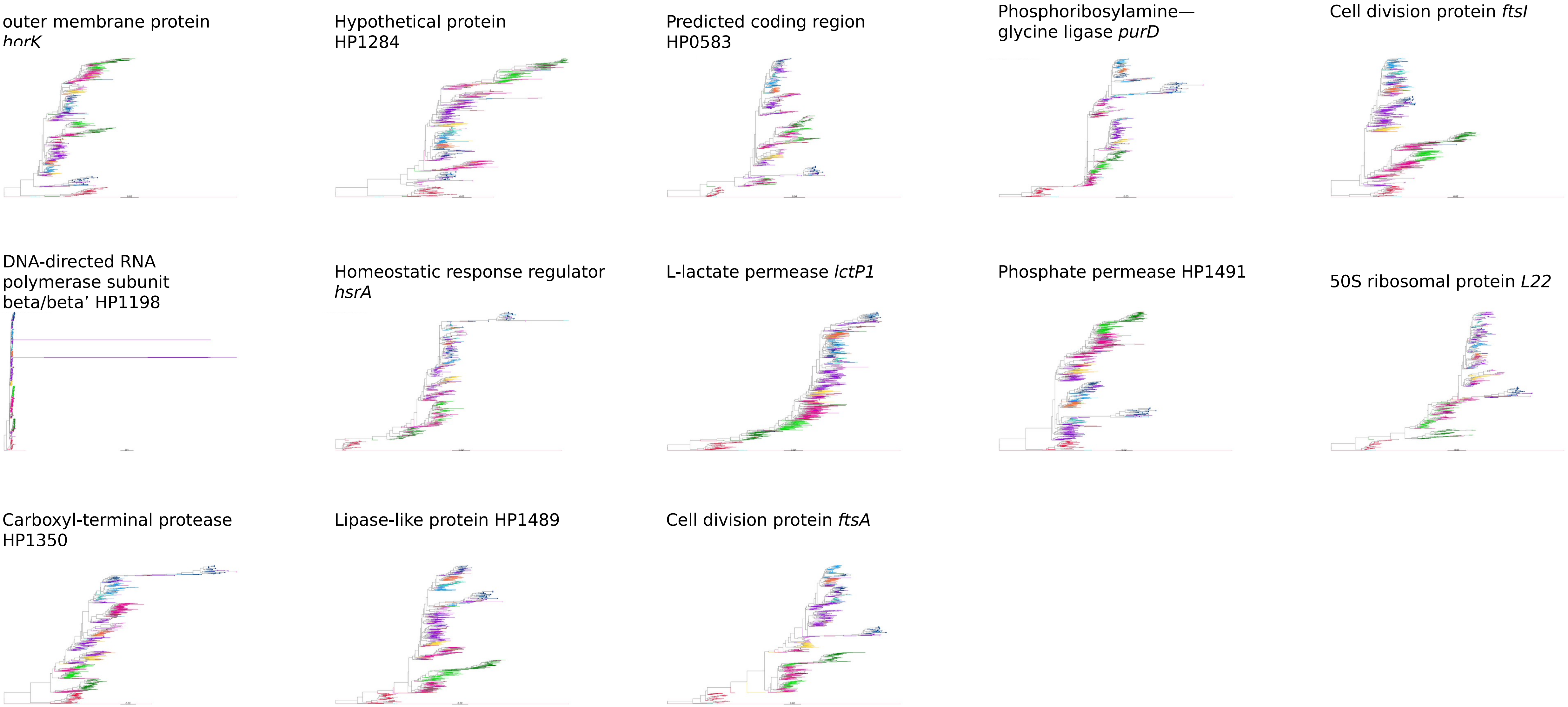

C

### Scenario 3: Presence of gene flow, in at least part of the gene

outer membrane protein
*horL*

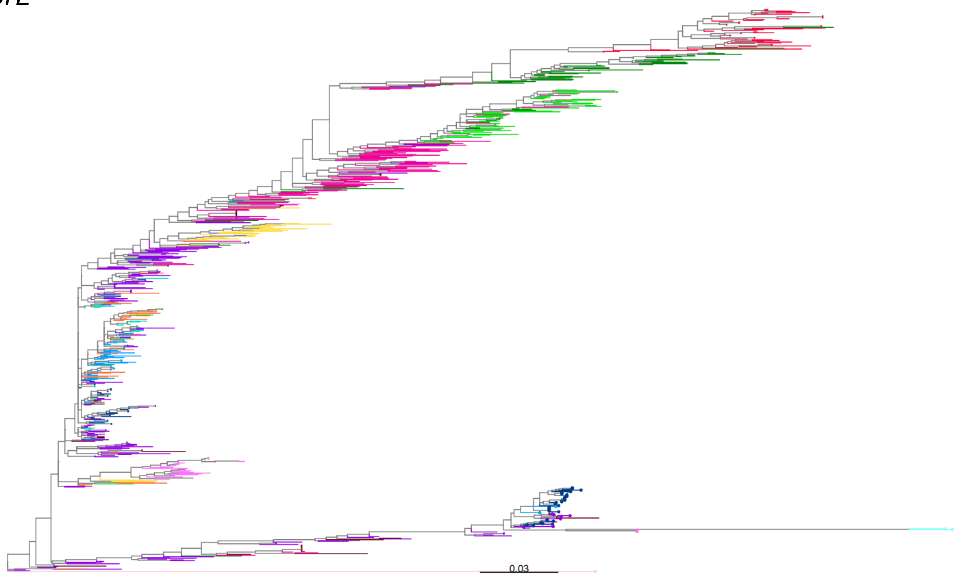


Methyl-accepting chemotaxis protein
HP0599

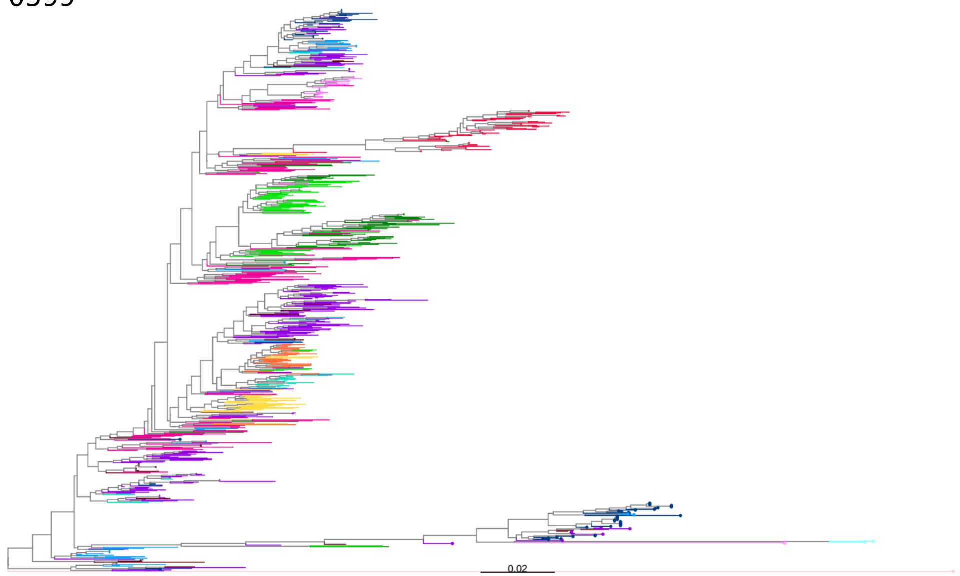


*lpp20* lipoprotein

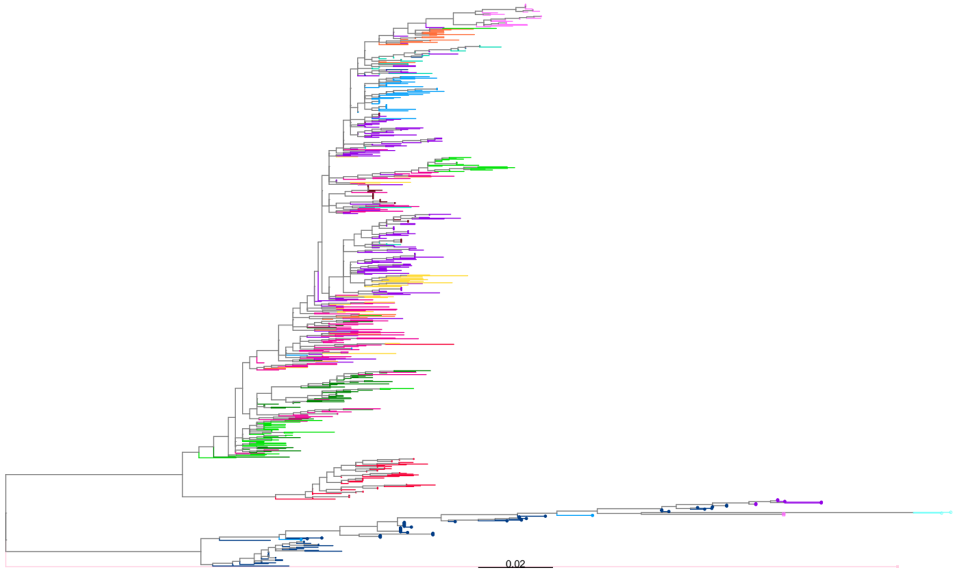

D Scenario 1bis: The ecospecies diverged after *H. cetorum* but before the hpAfrica2 branch split
Two copies are present

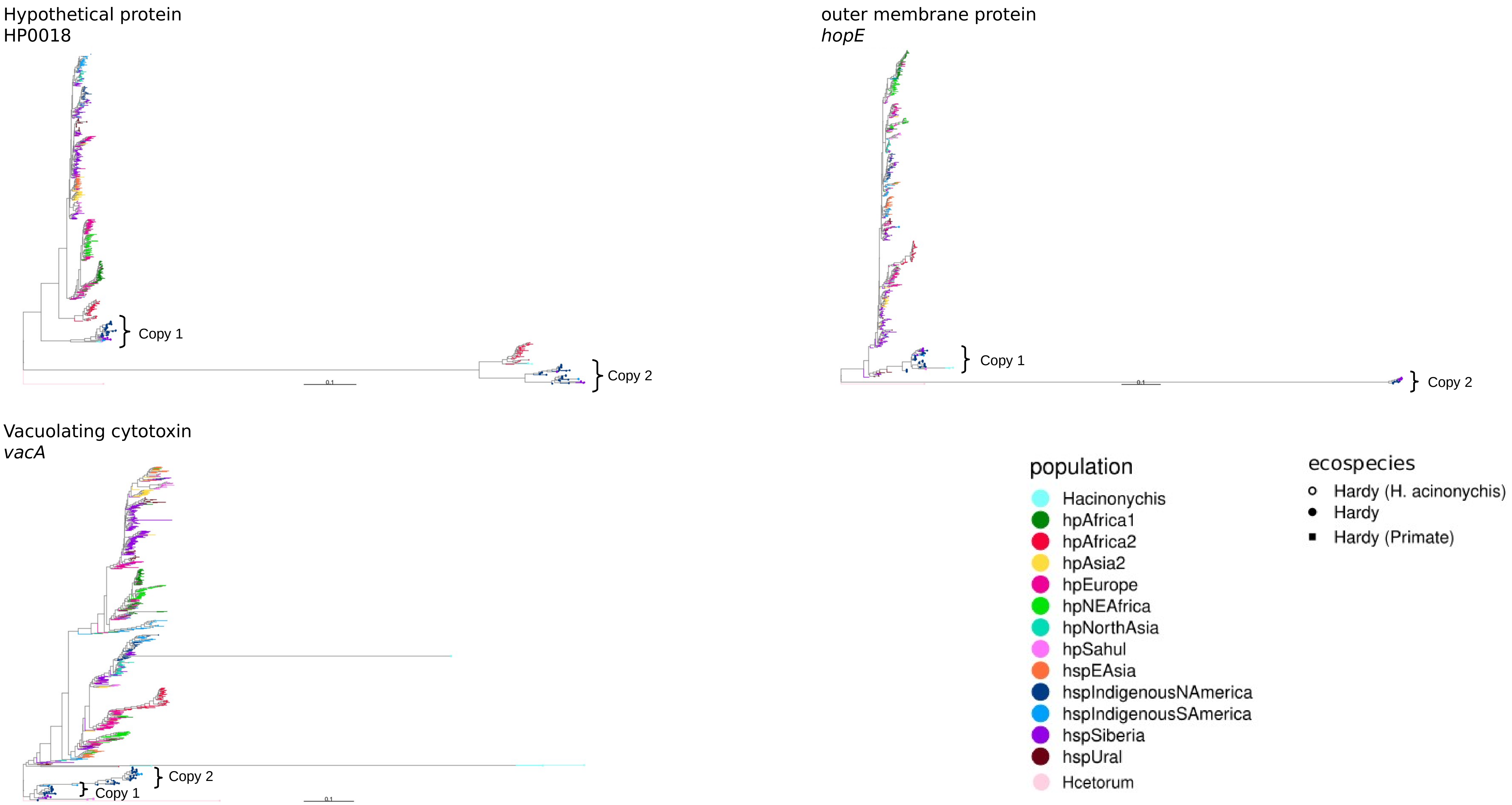

E

### Scenario 4: The polymorphism is older than *H. cetorum*

outer membrane protein
*hopF*

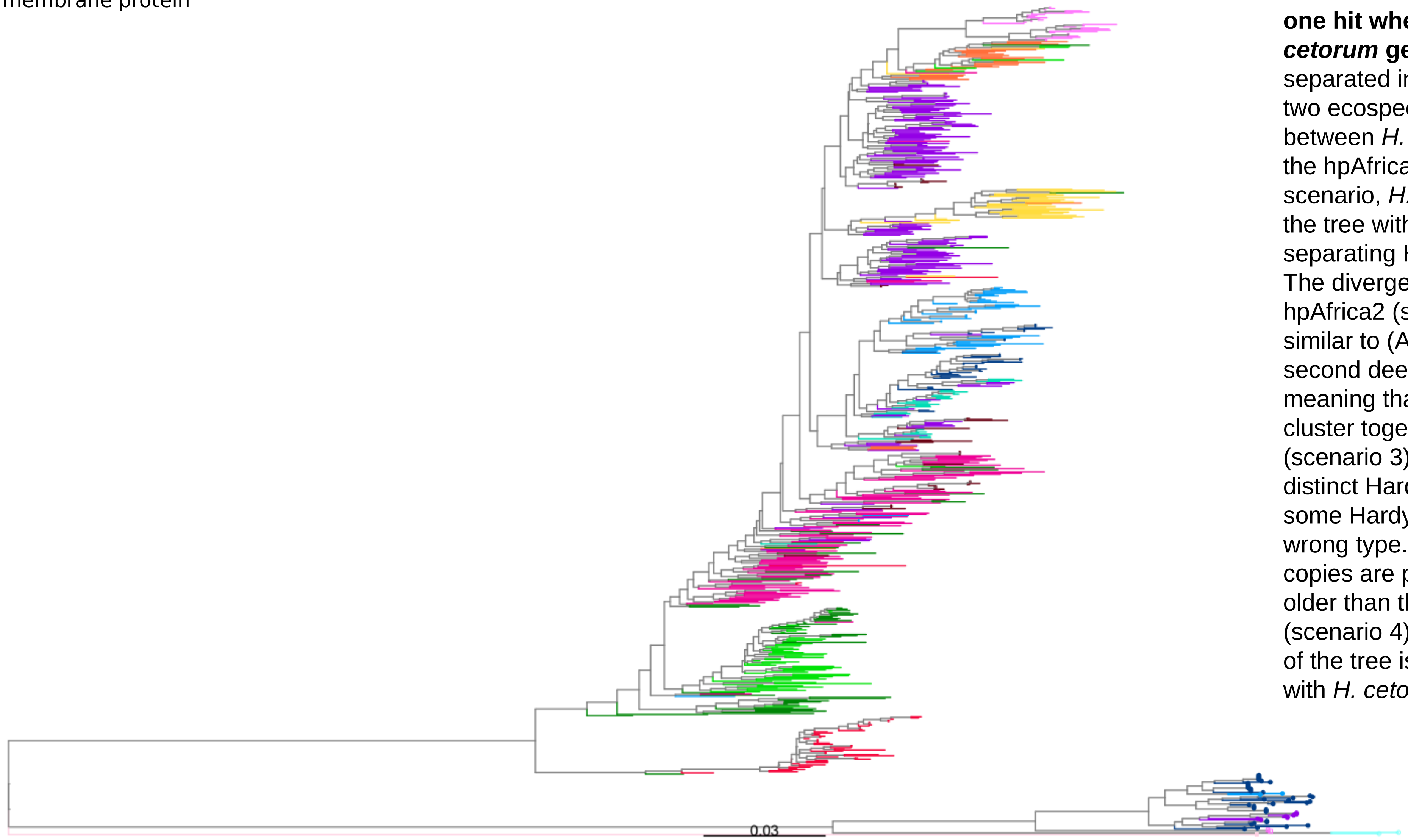


Figure S10:
**Phylogenetic trees of the sequences of differentiated genes that returned at least one hit when blasted against the *H. cetorum* genome.** The trees can be separated into four main scenarios: (A) The two ecospecies diverged after the divergence between *H. cetorum* and *H. pylori*, but before the hpAfrica2 branch split (scenario 1). In this scenario, *H. cetorum* is the deepest branch in the tree with the next deepest branch separating Hardy and Ubiquitous strains. (B) The divergence takes place after the split with hpAfrica2 (scenario 2). This scenario is similar to (A) except that *H. acinonychis* is the second deepest branch after *H. cetorum*, meaning that human and primate strains cluster together. (C) Presence of gene flow (scenario 3). In this scenario, there are distinct Hardy and Ubiquitous clusters but some Hardy or Ubiquitous strains have the wrong type. (D) Variant of scenario 1 but two copies are present. (E) The polymorphism is older than the divergence with *H. cetorum* (scenario 4). In this case, the deepest branch of the tree is between Hardy and Ubiquitous, with *H. cetorum* clustering with one or other.

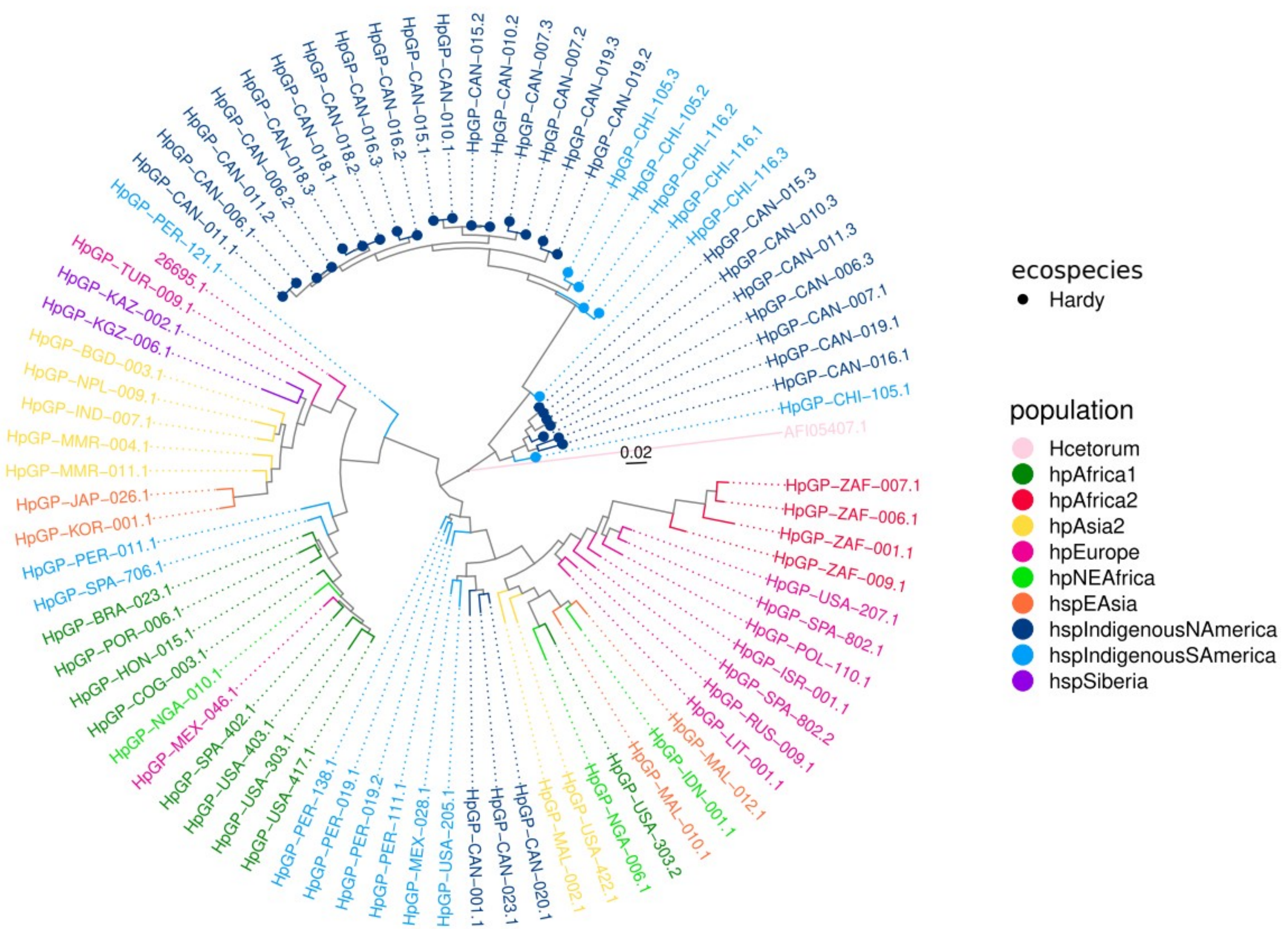


Figure S11:
**ML phylogenetic tree of *vacA*, based on nucleotide sequences from complete *H. pylori* genomes and *H. cetorum* (AFI05407.1).** The branches are colored based on their population and the Hardy strains are represented with a dot. Contrary to the *vacA* tree that is shown in Figure S10D, this tree was built using only complete genomes (in most of the non-complete genomes, *vacA* appears to be highly fragmented due to assembly issues). Only *one* copy of vacA from *H. cetorum* was included as they all clustered on the same branch. The leaves are labeled with strain name, plus the number of the copy of the gene in that strain (i.e., .1, and if there are more than one copy, .2 and .3). Note that, for the Hardy strains, the copies that are closer to the root (generally one copy per Hardy strain) are more *H. pylori*-like than the copies farther away.

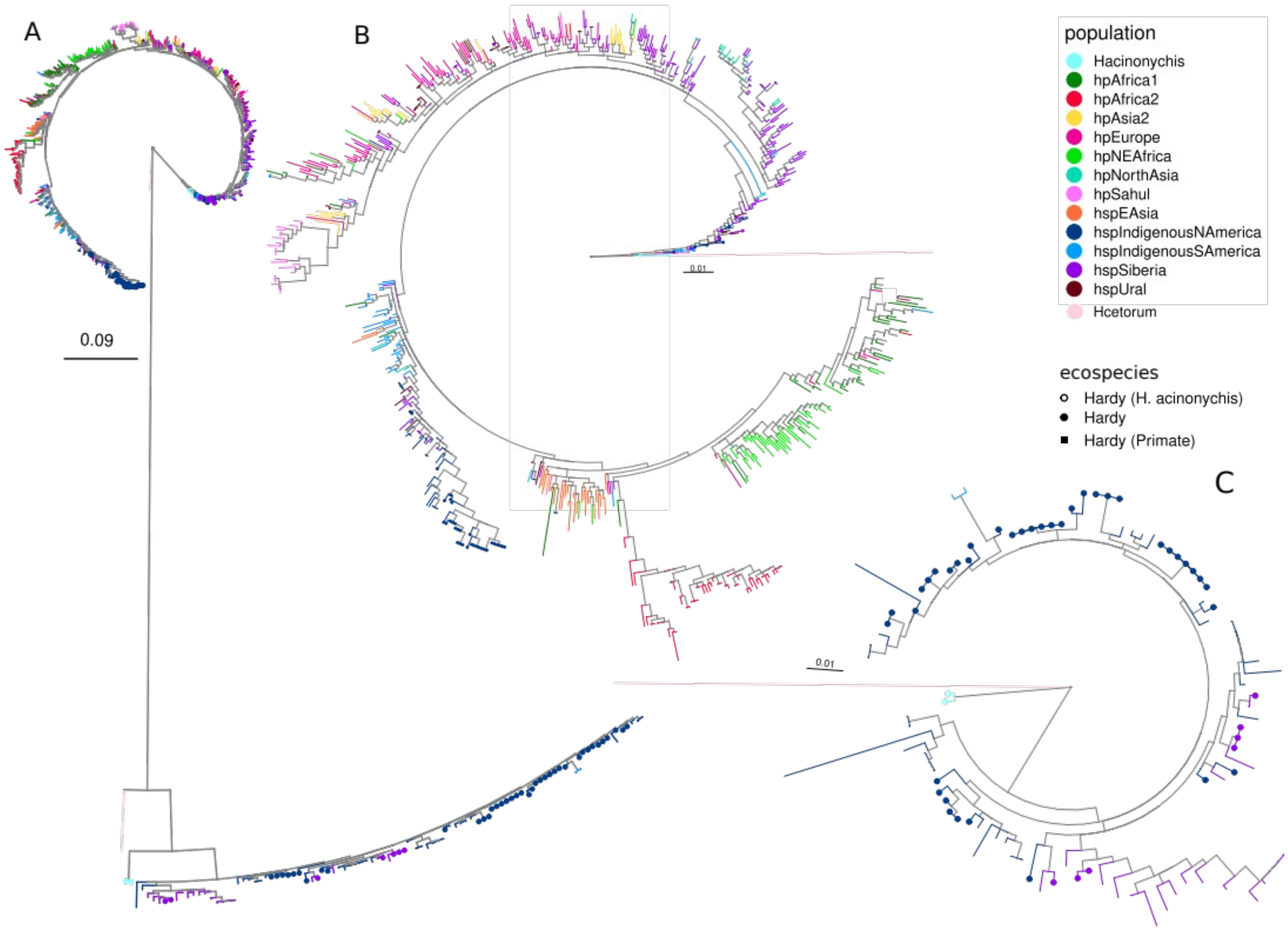


Figure S12:

**Phylogenetic tree of *ureA* sequences for a sample of strains.** (A) For all *ureA* copies, (B) for the first copy, which is conserved in all *H. pylori* strains and (C) for the second copy that is only present in the Hardy strains and in some of the Ubiquitous strains from the same populations as the Hardy ones. The strains are colored based on their population, a dot at the end of the branch indicates a Hardy strain. *H. cetorum* sequences are also included.

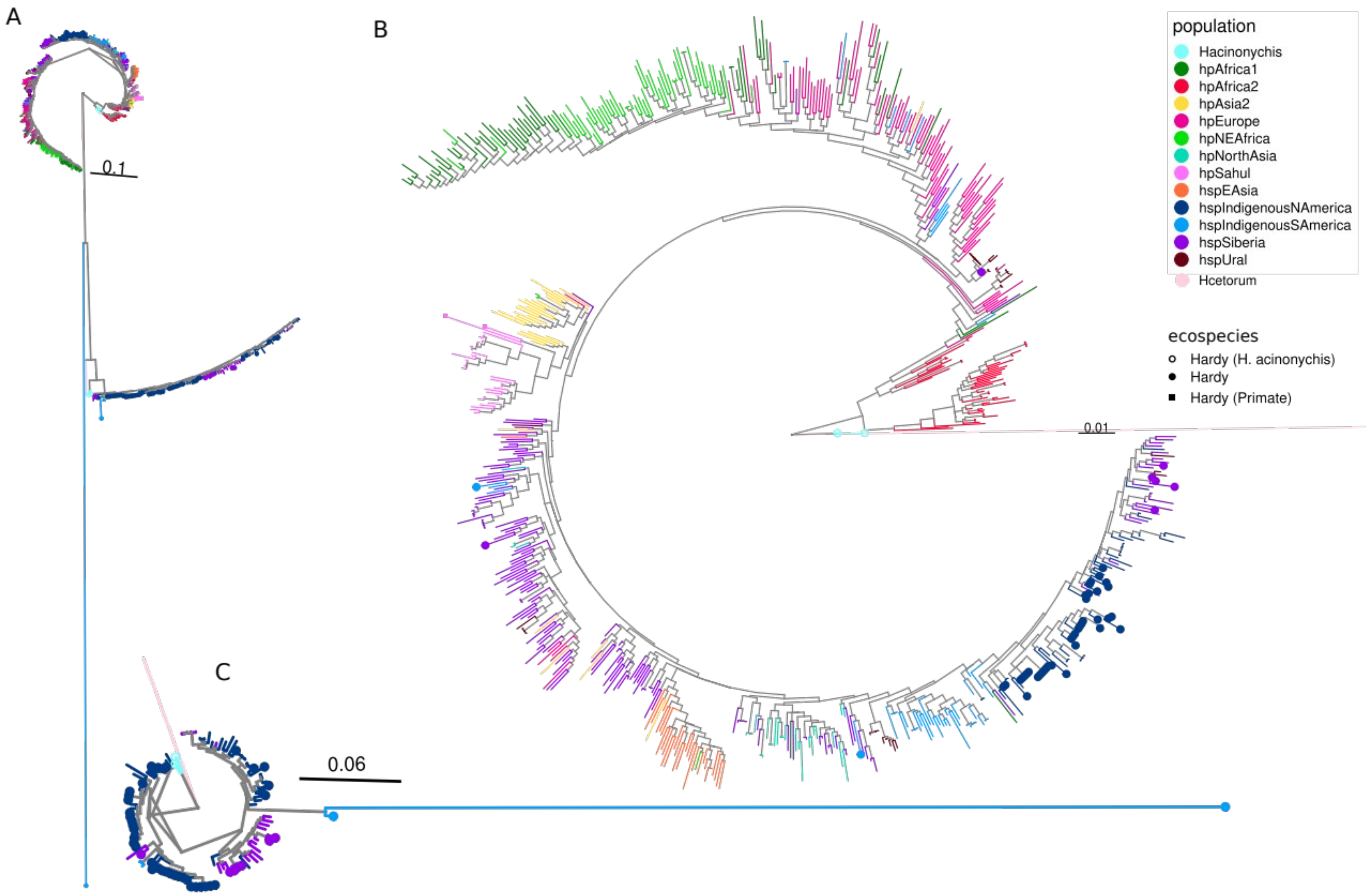


Figure S13:

**Phylogenetic tree of *ureB* sequences for a sample of strains.** (A) For all *ureB* copies, (B) for the first copy that is present in all strains and (C) for the second copy that is only present in the Hardy strains and in some of the Ubiquitous strains from the same populations as the Hardy ones. The strains are colored based on their population, a dot at the end of the branch indicates a Hardy strain. *H. cetorum* sequences are included.
